# *PMS1* as a target for splice modulation to prevent somatic CAG repeat expansion in Huntington’s disease

**DOI:** 10.1101/2023.07.25.550489

**Authors:** Zachariah L. McLean, Dadi Gao, Kevin Correia, Jennie C. L. Roy, Shota Shibata, Iris N. Farnum, Zoe Valdepenas-Mellor, Manasa Rapuru, Elisabetta Morini, Jayla Ruliera, Tammy Gillis, Diane Lucente, Benjamin P. Kleinstiver, Jong-Min Lee, Marcy E. MacDonald, Vanessa C. Wheeler, Ricardo Mouro Pinto, James F. Gusella

**Affiliations:** Molecular Neurogenetics Unit, Center for Genomic Medicine, Massachusetts General Hospital, Boston, MA 02114, USA; Department of Neurology, Harvard Medical School, Boston, MA 02115, USA; Medical and Population Genetics Program, the Broad Institute of M.I.T. and Harvard, Cambridge, MA 02142, USA; Center for Genomic Medicine and Department of Pathology, Massachusetts General Hospital, Boston, MA 02114, USA; Department of Pathology, Harvard Medical School, Boston, MA 02115, USA; Department of Genetics, Blavatnik Institute, Harvard Medical School, Boston, MA 02115, USA

**Author notes:** Materials & Correspondence: James F. Gusella.

## Abstract

Huntington’s disease (HD) is a dominantly inherited neurodegenerative disorder whose motor, cognitive, and behavioral manifestations are caused by an expanded, somatically unstable CAG repeat in the first exon of *HTT* that lengthens a polyglutamine tract in huntingtin. Genome-wide association studies (GWAS) have revealed DNA repair genes that influence the age-at-onset of HD and implicate somatic CAG repeat expansion as the primary driver of disease timing. To prevent the consequent neuronal damage, small molecule splice modulators (e.g., branaplam) that target *HTT* to reduce the levels of huntingtin are being investigated as potential HD therapeutics. We found that the effectiveness of the splice modulators can be influenced by genetic variants, both at *HTT* and other genes where they promote pseudoexon inclusion. Surprisingly, in a novel hTERT-immortalized retinal pigment epithelial cell (RPE1) model for assessing CAG repeat instability, these drugs also reduced the rate of *HTT* CAG expansion. We determined that the splice modulators also affect the expression of the mismatch repair gene *PMS1*, a known modifier of HD age-at-onset. Genome editing at specific *HTT* and *PMS1* sequences using CRISPR-Cas9 nuclease confirmed that branaplam suppresses CAG expansion by promoting the inclusion of a pseudoexon in *PMS1*, making splice modulation of *PMS1* a potential strategy for delaying HD onset. Comparison with another splice modulator, risdiplam, suggests that other genes affected by these splice modulators also influence CAG instability and might provide additional therapeutic targets.

## Introduction

Huntington’s disease (HD, MIM: 143100) is a dominantly inherited neurodegenerative disorder whose motor, cognitive, and behavioral manifestations are caused by an expanded CAG repeat in the first exon of *HTT* ^1^, which encodes huntingtin. The inherited repeat, whose length is negatively correlated with HD age-at-onset, undergoes further expansion in various somatic tissues but particularly in the brain ^2,3^, with the largest post-mortem expansions found in those individuals with the earliest onset ^4^. Human genome-wide association studies (GWAS) have revealed that HD age-at-onset is influenced by some DNA repair genes that play a role in repeat instability ^5^. These feature, together with the similar age-at-onset and onset lack of increased severity in individuals with two expanded alleles ^6,7^, have led to a sequential two-step model for HD pathogenesis wherein 1) the inherited CAG repeat lengthens over an individual’s life in cells that enable CAG repeat expansion, and 2) once the CAG repeat reaches a cell type-specific threshold length, it triggers toxicity/dysfunction that leads eventually to cell death ^8,9^. The ultimate mechanism of toxicity is still unclear. Candidates include dysfunction caused by mutant huntingtin or amino-terminal fragments containing a lengthened CAG repeat-encoded polyglutamine segment ^10^ and toxicity via *HTT* mRNA ^11,12^.

The two-step mechanism proposed to explain HD pathogenesis also suggests two distinct therapeutic options, one to prevent CAG repeat expansion by early intervention and the other to reduce the toxicity process initiated by the somatically expanded CAG repeat. To date, more translational attention has been paid to the toxicity step, where attempts to reduce *HTT* mRNA/protein level by targeted genetic approaches have included antisense oligonucleotides (ASOs) and RNA interference, and *HTT* transcript splice modulation ^13^. Branaplam (Novartis) and PTC518 (PTC Therapeutics) are small molecules that have been in phase II clinical trials for HD based on their modulation of *HTT* splicing. The chemical structure of PTC518 has not been disclosed, but PTC Therapeutics has previously reported that risdiplam, a drug used for the treatment of spinal muscular atrophy (SMA), also targets *HTT* with lower potency ^14^. These splice modulators stabilize non-canonical nGA 3’-exonic motifs, resulting in the inclusion of a frame-shifting pseudoexon between *HTT* exons 49 and 50 ^14,15^, with consequent lowering of huntingtin level. Recently, the VIBRANT-HD clinical trial of branaplam in adults with HD (phase 2b, Novartis) clinical trial was halted due to safety concerns, highlighting the need for further research into the effect on HD cells, including the role of off-target splice modulation.

For designer therapeutics based on genetic targets, polymorphic sequence variation can potentially affect both on- and off-target efficacy. Consequently, we explored the effects of genetic variation surrounding the *HTT* pseudoexon and predicted alternative targets in other loci in human lymphoblast cell lines (LCLs) of defined genotype. We found that the effectiveness of the splice modulators branaplam and risdiplam can be influenced by genetic variants, both at *HTT* and other genes where they promote pseudoexon inclusion. Interestingly, these drugs also reduced the rate of *HTT* CAG expansion in a novel *in vitro* model of repeat instability. We show the splice modulators also target *PMS1*, a known modifier of HD age-at-onset, and demonstrate that branaplam’s suppression of CAG expansion is due to pseudoexon inclusion in *PMS1*, making this a potential strategy for treatment of HD.

## Results

### Splice modulator-induced products and dose-response

We treated lymphoblastoid cell lines (LCLs) from HD individuals with branaplam or risdiplam to confirm splice modulation of *HTT*. In each case, two alternatively spliced products were produced. One RNA included the pseudoexon (exon 50a in Figure 1a,b) from novel 3’ and 5’ splice sites (ss) between the exon 49 and exon 50 sequences. The other resulted in the lengthening of exon 50 (exon 50b) via the use of the same novel alternative 3’ss as the pseudoexon (Figure 1a,b, Supplementary Figure 1). Both alternatively spliced products are predicted to share the same functional outcome since the inclusion of these pseudoexons introduces a premature termination codon into the *HTT* transcript. The compounds produced a dose-dependent decrease in the *HTT* canonical isoform (Figure 1c), with branaplam (IC50 25 nM) approximately 25 times more potent than risdiplam (IC50 636 nM), but they differed in the relative proportion of the two novel products. Branaplam produced a mean ratio of exon 50b to exon 50a of 2.7 across the concentration gradient, while risdiplam displayed a lower exon 50b to exon 50a ratio of 0.30 (Figure 1c).

**Figure 1.**
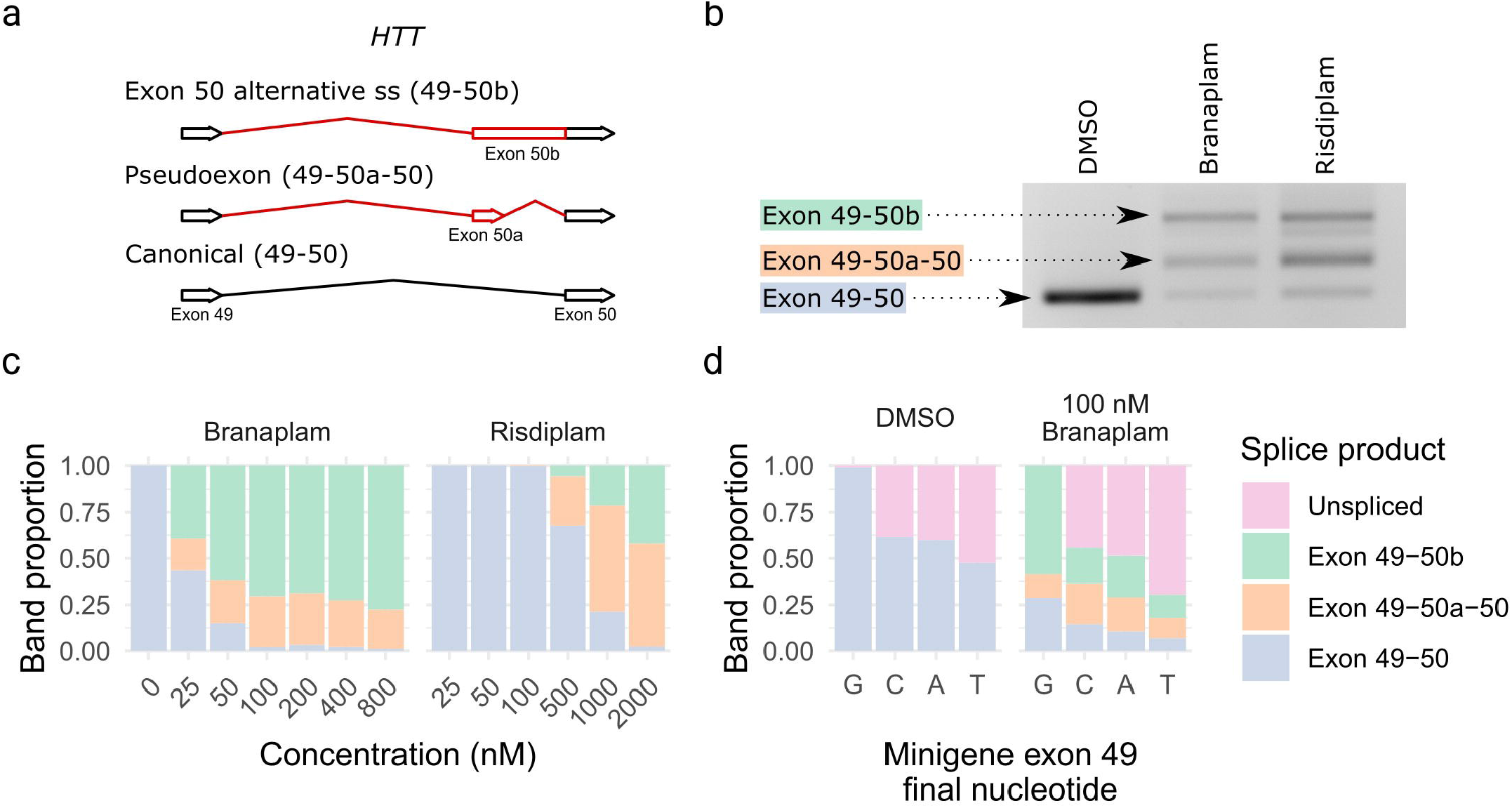
Branaplam and risdiplam treatment of HD LCLs produced two major *HTT* alternative splice products. (a) Schematic diagram showing the alternative *HTT* splice products upon drug treatment. (b) PCR from exon 49-50 showing the size of the splice products. (c) Branaplam and risdiplam dose response for each *HTT* splice product. (d) Quantification of splice products produced from mutant minigenes following transfection of HEK 293T cells either treated with a vehicle control (DMSO) or 100 nM branaplam.

Branaplam has been shown to bind the novel exon 50a 5’ss with U1 snRNP to enable the formation of this pseudoexon ^14^. Therefore, the prominent production of a new product in which pseudoexon 50b is generated by the novel 3’ss but utilizes the canonical exon 50 5’ss was unexpected. We postulated that generation of the exon 50b product might be influenced by the relative strength of neighboring splice site strengths. We reasoned that, due to the stronger upstream exon 49 5’ss, the initial portion of intron 49 up to the pseudoexon 50a 3’ss might be spliced out first, but the intron section downstream of the pseudoexon 50a 5’ss be retained due to the relative weakness of the latter. This hypothesis predicted that weakening the upstream site would decrease the exon 50b/exon 50a ratio produced after drug treatment. Therefore, we used site-directed mutagenesis in a minigene construct to vary the final base of exon 49 from the normal GAG|gt exon-intron junction (highlighted by “|”) to GA<u>C</u>|gt, GA<u>T</u>|gt, and GA<u>A</u>|gt (mutated nucleotide underlined). When transfected into HEK293T cells and analyzed with PCR specific to the minigene, without branaplam, the GA<u>C</u>|gt and GA<u>A</u>|gt mutants each resulted in ∼30% unspliced minigene product, with GA<u>T</u>|gt at 50% unspliced. With branaplam treatment, the ratio of exon 50b/exon 50a decreased from 4.6 for the GAG|gt minigene to 0.9 for GA<u>C</u>|gt, 1.1 for GA<u>T</u>|gt, 1.2 for GA<u>A</u>|gt (Figure 1d), indicating that the relative strength of the upstream exon 49 5’ss influences branaplam-induced splicing outcomes.

### Rare sequence variants affect HTT splice modulation

Given this evidence for sequence context having an impact on the effects of branaplam treatment, we evaluated the effect of genetic variation surrounding the *HTT* pseudoexon on drug-induced splice modulation. Population-based estimates from gnomAD (global ancestry) indicated that *HTT* intron 49 has low genetic variation, with no variants of minor allele frequency (MAF) > 10% and only two > 1% (Figure 2a). We screened our bank of previously genotyped HD LCLs and identified 15 lines collectively representing eight single nucleotide variants (SNVs) of interest. We included one common variant (rs362331) located in exon 50 (Figure 2a) and seven less frequent variants distributed across intron 49, with two close to the 5’ss (rs193157701, rs79689511), one close to the intron 49 3’ss (rs376150131), and four centrally located (rs10030079, rs145498084, rs567263187, rs772437678). Of the latter, rs772437678 and rs145498084 are located 11 and 21 nucleotides upstream of the pseudoexon 3’ss, respectively (Figure 2a). We did not have cell lines with rs148430407, a rare SNV located 2 nucleotides downstream of the pseudoexon 5’ss that alters the canonical 5’ss intron sequence from gt to g<u>g,</u> primarily in individuals of African ancestry.

**Figure 2.**
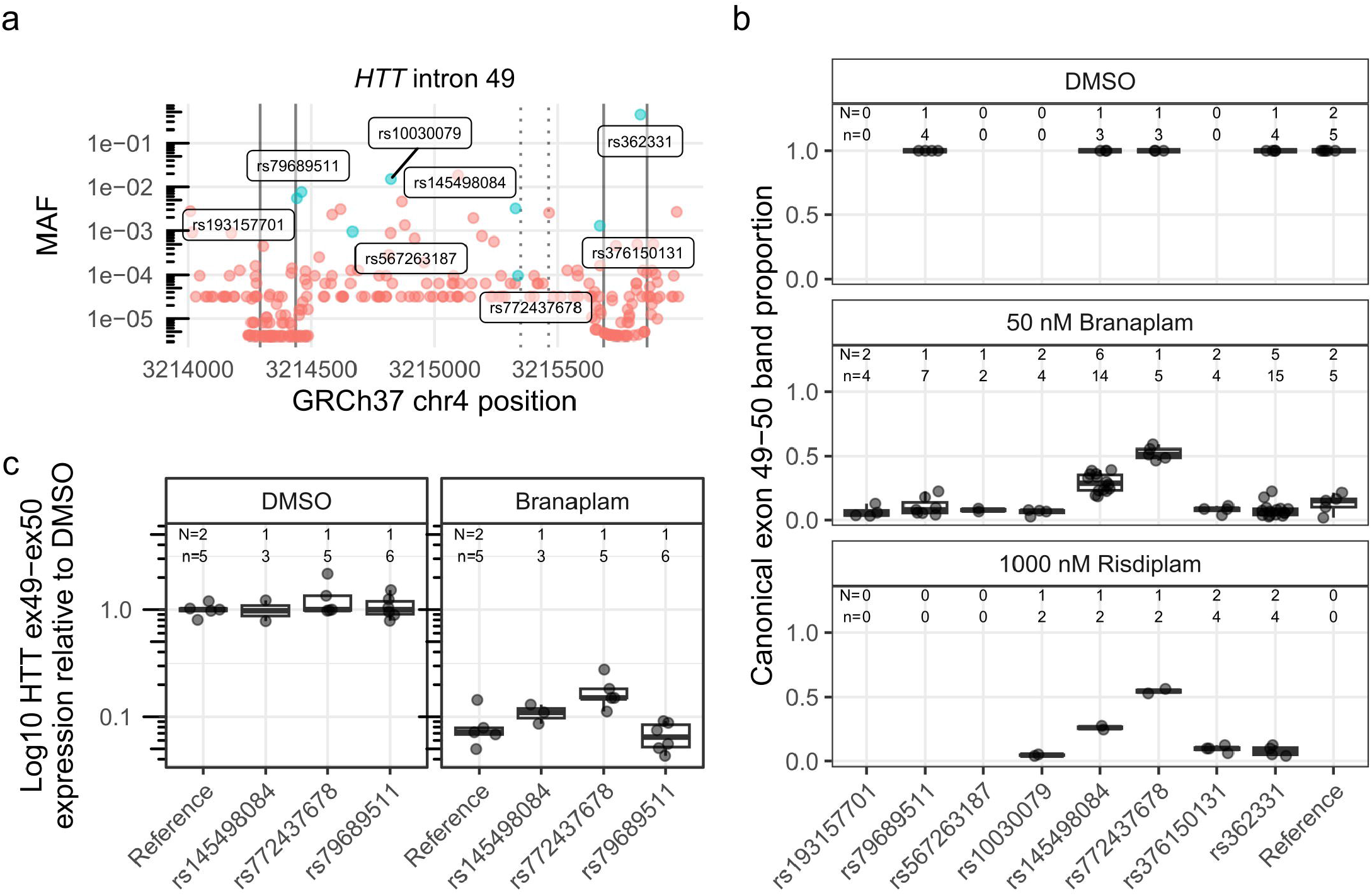
Two single nucleotide variants affected *HTT* splice modulation. (a) Minor allele frequency (MAF) of variants spanning *HTT* exon 49-50 (exons marked with solid vertical lines), with variants represented in the cell lines tested labeled and highlighted in blue. The dotted vertical lines indicate the pseudoexon splice sites (ss). (b) The proportion of canonical *HTT* exon 49-50 product across tested cells lines, grouped by heterozygous presence of variant. Since the production of the pseudoexon requires drug treatment, only a subset of the cell lines were treated with DMSO control. (c) Absolute quantification by ddPCR across exon 49-50 junction for a subset of the cell lines on a log10 axis. N = Number of cell clones, n = cultures analyzed

Treatment of the HD LCLs with 50 nM branaplam reduced the proportion of canonical splice product to 0.098 (95% CI: 0.021 to 0.17) in cells homozygous for the reference sequence, but only to 0.49 (95% CI: 0.41 to 0.57) and 0.32 (95% CI: 0.25 0.39) (p < 0.0001 in both cases) in cells heterozygous for rs772437678 or rs145498084, respectively (Figure 2b). The remaining cell lines with variants of interest showed a similar proportion of canonical splice product to those with the reference sequence (p ≥ 0.2). The relatively higher fraction of canonical splice product remaining in cell lines with rs772437678 and rs145498084 is presumed to derive from interference by the minor allele of the respective SNV with the branaplam mechanism. We observed a similar result for these two SNVs with 1000 nM risdiplam treatment (Figure 2b). Of note, the common SNV rs362331, located 28 bases upstream of the exon 50 5’ss, did not affect splice modulation (p > 0.17). Given the robust interference with splice modulation by rs772437678, we repeated the branaplam dose-response experiment with cell lines respectively heterozygous for rs772437678 or, as a control, rs79689511. At higher branaplam concentrations, the proportion of the canonical isoform continued to decrease in cell lines with rs772437678 but was consistently higher than in the control (Supplementary Figure 2). We also analyzed the exon 50b/exon 50a ratio for this set of cell lines and observed no differences from the samples with reference sequence (p ≥ 0.1) (data not shown).

Although the densitometric method permits comparison of the relative levels of canonical and non-canonical splice variants, we expected that the absolute level of *HTT* mRNA might be reduced by preferential degradation through nonsense-mediated mRNA decay (NMD) of the non-canonical products due to their premature termination codon. Consequently, we performed droplet digital PCR (ddPCR) for accurately quantifying the *HTT* canonical isoform, analyzing a subset of the same samples using a hydrolysis probe spanning the exon 49-50 junction. Treatment with 50 nM branaplam reduced *HTT* cDNA with the exon 49-50 junction by ∼15-fold in control cells and ∼7-fold in the cell line with rs772437678, reflecting the ∼2-fold relative effect seen by densitometry (Figure 2c).

### SpliceAI predictions on branaplam-responsive exons genome-wide

Given that sequence variants near the splice site altered the effect of the splice modulators at the intended target locus, we next used the deep neural network tool SpliceAI ^16^ to predict variants that might modulate branaplam-responsive exons from off-target genes identified transcriptome-wide from previously published datasets: Monteys et al. 2021 ^17^ (HEK293, 25 nM Branaplam); Bhattacharyya et al., 2021 ^14^ (SH-SY5Y cells, 100 nM Branaplam); Keller et al., 2022 ^15^ (SH-SY5Y cells, 100 nM branaplam) (Figure 3a). From the combined set of pseudoexons (Supplementary data 1), SpliceAI identified primarily rare variants within the 50 base pairs (bp) adjacent to pseudoexon splice junctions (Figure 3b, Supplementary data 2). Near the *HTT* pseudoexon, only rs772437678, which interferes with the branaplam effect, and rs148430407, which we were unable to test, yield significant negative SpliceAI scores, consistent with a reduction in pseudoexon inclusion. At MAF >1 %, single variants in other genes showed significant SpliceAI scores. Four of these variants are predicted to enhance the incorporation of a pseudoexon in two genes (positive SpliceAI score), sensitizing *DLGAP4 and ZNF680* to the splice modulation, while variants in *ATF6*, *ENOX1* and *TENT2* are predicted to interfere with pseudoexon inclusion (negative SpliceAI score). SpliceAI also predicted six frequent variants (MAF >1%; three positive and three negative SpliceAI scores) in five genes among those reported to display branaplam-responsive alternative splicing of annotated exons.

**Figure 3.**
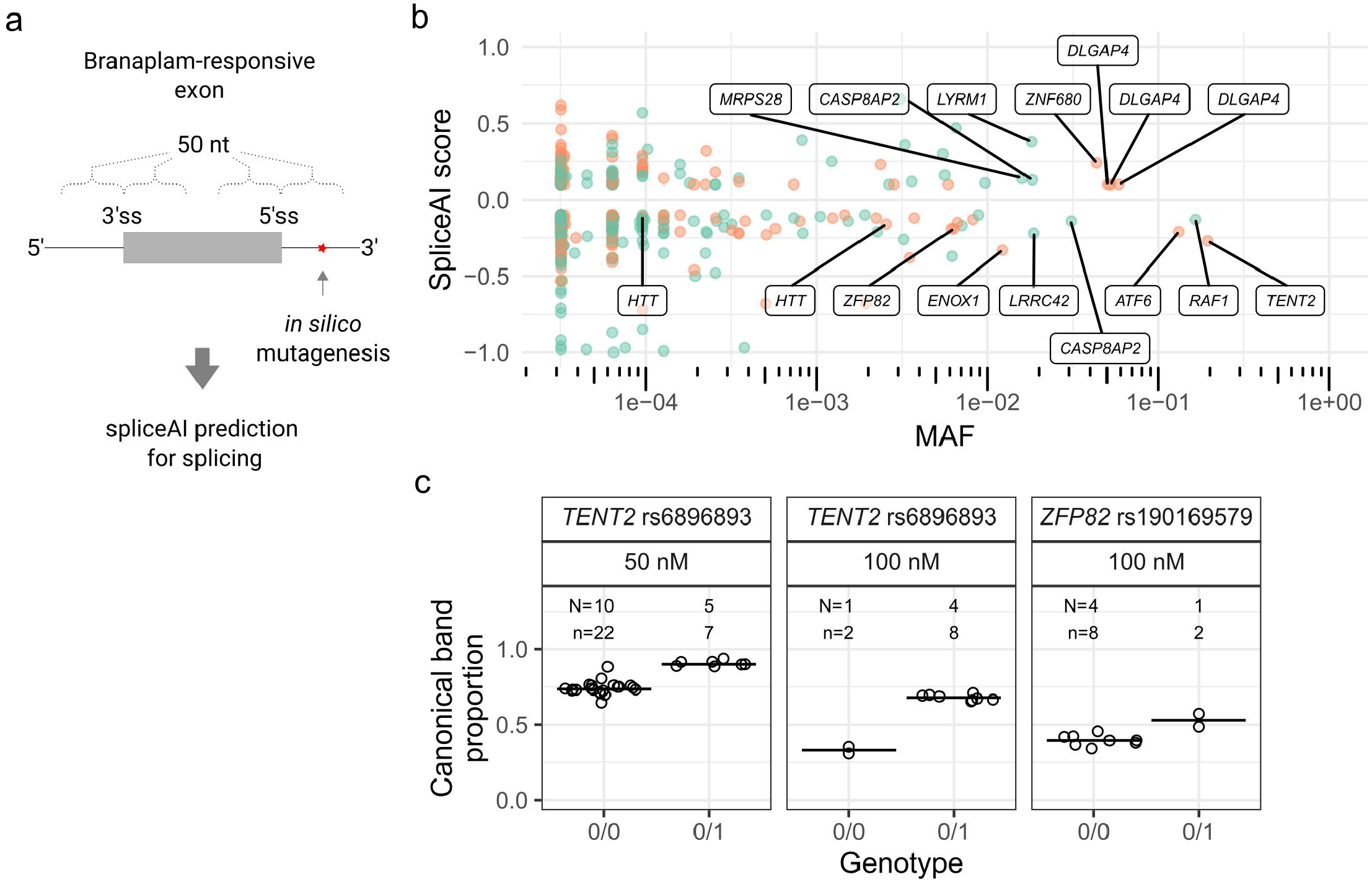
SpliceAI identified variants predicted to affect splicing of genome-wide branaplam- responsive exons. (a) SpliceAI predictions were made for variants within 50 nt of branaplam- responsive exon and pseudoexon splice junctions. (b) Variants near branaplam-responsive pseudoexons (orange) and exons (green) that yield significant SpliceAI scores are plotted by allele frequency with gene names indicated for selected variants. *HTT* variants rs148430407 (MAF 2.6x10^-3^) and rs772437678 (MAF 9.6x10^-5^) are labelled, while rs145498084 did not have a significant SpliceAI score (c) SpliceAI-predicted variants affect splice modulation of *TENT2* and *ZFP82*. Proportion of canonically spliced product across tested LCLs for *TENT2* and *ZFP82*, grouped by no presence (0/0) or heterozygous presence (0/1) of variant. N = Number of cell lines for variant, n = cultures analyzed.

We validated the SpliceAI results with two variants predicted to have a negative effect on pseudoexon splicing probability in *TENT2*, (rs6896893, spliceAI score -0.27, MAF 19%) and *ZFP82* (rs190169579, spliceAI score -0.19, MAF 0.63%), respectively. First, we confirmed that branaplam treatment results in pseudoexon inclusion for both genes (Supplementary Figure 3b). When treated with 50 nM branaplam, LCLs heterozygous for the *TENT2* SNV showed less pseudoexon inclusion (i.e., a higher proportion of canonical transcript) than those homozygous for the major allele (0.91, 95% CI: 0.88 to 0.93 versus 0.74, 95% CI: 0.73 to 0.76; p < 0.0001) (Figure 3c). Treatment with 100 nM branaplam further accentuated this effect (0.68, 95% CI: 0.66 0.69, versus 0.33, 95% CI: 0.30 to 0.36; p = 0.0002) (Figure 3c). Similarly, LCLs heterozygous for the *ZFP82* SNV treated with 100 nM branaplam showed a higher proportion of canonical *ZFP82* transcript, 0.53 (95% CI: 0.47 to 0.58) compared to 0.40 (95% CI: 0.37 to 0.42) (p = 0.02) in LCLs without the minor allele (Figure 3c).

### Branaplam and risdiplam suppress CAG repeat expansion

Having established that consideration of DNA sequence polymorphisms can be relevant to both the proposed HD therapeutic’s on-target and off-target splicing effects, we turned our attention to the critical driver of HD pathogenesis: *HTT* CAG repeat expansion. It has been suggested that reducing huntingtin levels by ASO treatment also reduces CAG repeat expansion ^18^. Therefore, we tested the effects of the *HTT*-lowering splice modulators on CAG repeat expansion. Most cultured HD cell lines display limited CAG repeat instability, so we developed a new model system for this purpose in hTERT-RPE1 (RPE1) cells. RPE1 is a near-diploid immortalized cell line often used to study DNA repair pathways ^19^. It can be arrested at G0/1 through contact inhibition by growing the cells to confluency. We isolated the expanded CAG *HTT* exon 1 from a juvenile-onset HD individual (115 CAGs) and knocked the fragment into the AAVS1 safe harbor locus (intron 1 of *PPP1R12C* on chromosome 19) under a doxycycline-inducible promoter, intending to control transcription and transcription-linked repeat instability. We isolated 8 clones, each with 110-115 CAG repeats, and cultured the cells in the presence and absence of doxycycline. The non-induced lines showed relatively rapid CAG repeat expansion with an average CAG weekly gain of 0.87 units (95% CI: 0.74 to 1.0). In the presence of doxycycline, the lines showed much less repeat expansion with only 0.051 (95% CI: -0.082 to 0.18) CAG gain per week (p < 0.0001) (Figure 4a, b).

**Figure 4.**
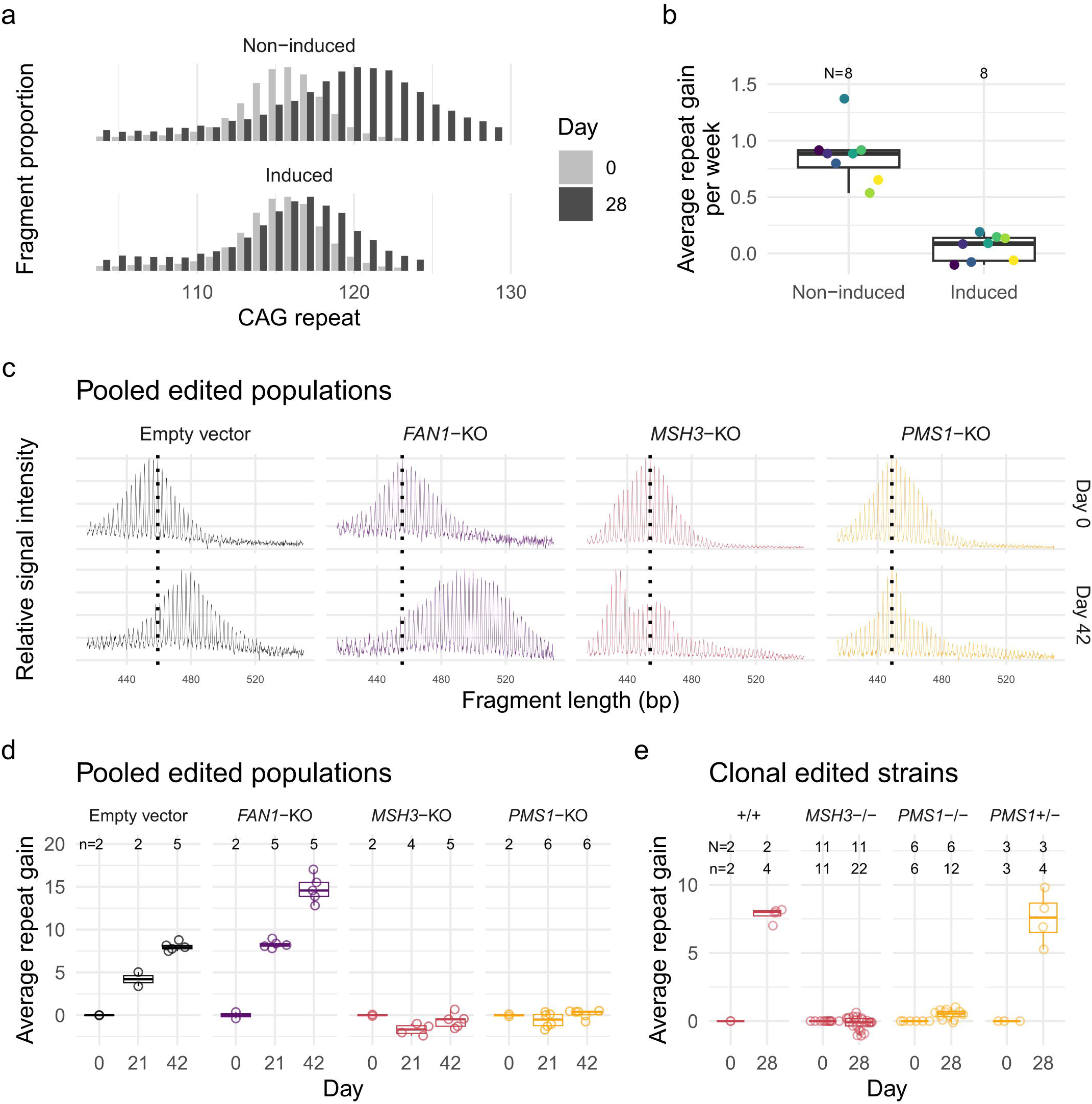
RPE1-AAVS1-CAG115 cell model for CAG repeat instability. (a) CAG repeat fragment distribution for a single RPE1-AAVS1-CAG115 clone in the absence (top) or presence (bottom) of doxycycline-induced transcription either at day 0 (light gray) or 28 (dark gray). (b) The average repeat gain per week for the 8 RPE1-AAVS1-CAG115 clones with either non- induced or induced transcription. Color indicates cell clone and N the total number of clones analyzed (c) Fragment analysis traces showing the change in CAG repeat length distribution across time in different non-edited and edited cells for pooled edited populations. Color indicates CRISPR-Cas9 target: non-targeting empty vector (black), *FAN1* (purple), *MSH3* (red), and *PMS1* (orange). The plots represent raw fluorescent signal without baseline correction and therefore have a negative signal bias with increasing fragment size. The following instability metrics were derived from data processed in the GeneMapper software which corrects this bias. (d) Average repeat gain for pooled edited populations, with each dot representing a biological replicate. (e) Average repeat gain for cell clones isolated from either *MSH3* (red) or *PMS1* (orange) targeted populations. N = Number of cell clones, n = cultures analyzed.

We validated the relevance of our RPE1-AAVS1-CAG115 cell line to model somatic instability processes by perturbing modifiers of HD age-at-onset predicted to influence repeat instability ^9^. We utilized CRISPR-Cas9 nuclease to target and modify the coding sequences of *FAN1*, *MSH3*, and *PMS1* via loss-of-function insertion or deletion mutations (indels) and analyzed repeat instability in the pooled heterogeneously-edited populations of cells (Supplementary Figure 4a). Fragment analysis traces for the empty vector control and *FAN1-*targeting vector each showed a single approximately normally distributed population increasing in CAG length over time (Figure 4c). As expected, *FAN1* knockout increased the average CAG repeat gain per week from 1.34 (95% CI: 1.22-1.47) to 2.52 (95% CI: 2.40-2.64) (p < 0.0001) (Figure 4d). By contrast, *MSH3* and *PMS1* knockouts produced more complex distributions (Figure 4c). The *MSH3* knockout culture developed a clear bimodal CAG repeat length distribution with one peak appearing to reflect CAG repeat contraction and the other modest, if any, expansion. The *PMS1* knockout exhibited a small degree of expansion in some cells, albeit far less than that seen in either the empty vector or *FAN1* knockout conditions.

To clarify the different instability distributions for contractions versus expansions in *MSH3*- and *PMS1*-edited cells, we isolated clones from the pooled populations and repeated the instability analysis. For each of the genotypes, the distribution was monomodal (Supplementary figure 4b), suggesting that the above distributions reflected a mixture of edited and non-edited cells that differed in their propensity for CAG expansion. From the *MSH3*-targeted population, we obtained 3 non-edited and 11 biallelically-edited clones representing complete knockouts. The latter showed an average repeat loss of 0.037 (95% CI: -0.11 to 0.035) per week compared to a gain of 2.0 (95% CI: 1.8 to 2.1) for the non-edited lines (Figure 4e). For *PMS1*, we derived 3 monoallelically-edited and 6 biallelically-edited clones. The heterozygous *PMS1* lines did not differ (p = 0.63) from non-edited cells with a repeat gain of 1.9 (95% CI: 1.7 to 2.1) per week.

For the biallelically edited strains, there was a small amount of repeat expansion with a repeat gain of 0.13 (95% CI: 0.028 to 0.22) per week, which was significantly higher (p = 0.0086) than the equivalent *MSH3* knockouts. The *PMS1* genome editing was in exon 6, which can be alternatively spliced, so the residual repeat expansion might be due to expression of a minor isoform in RPE1 cells (Supplementary figure 8a). Overall, these results are consistent with the effects of these HD genetic modifiers in HD individuals and animal and other cell models. The rapid CAG expansion in this system makes the RPE1-AAVS1-CAG115 a useful model for functional genomic investigations of CAG repeat instability.

We next used an individual RPE1-AAVS1-CAG115 clone, maintained at confluency without doxycycline, to quantify repeat instability for experiments with high or low dosages of branaplam or risdiplam as a test for an effect on CAG repeat expansion. Branaplam caused a dose-dependent reduction in repeat expansion, with an average CAG gain per week of 0.94 (CI:0.88- 1.01) in the DMSO control, 0.81 (CI:0.75-0.88) at 25nM branaplam (p = 0.005) and 0.73 (CI:0.66-0.79) at 100 nM branaplam (p < 0.0001) (Figure 5a). By contrast, 100 nM risdiplam produced relatively little change in CAG repeat gain, at 0.87 (CI:0.81-0.93) per week compared to the control’s 0.77 (CI:0.71-0.83) (p = 0.03). However, 500 nM risdiplam caused a significant decrease in the rate of repeat expansion to 0.40 (CI:0.34-0.46) CAGs per week (p < 0.0001) (Figure 5b).

**Figure 5.**
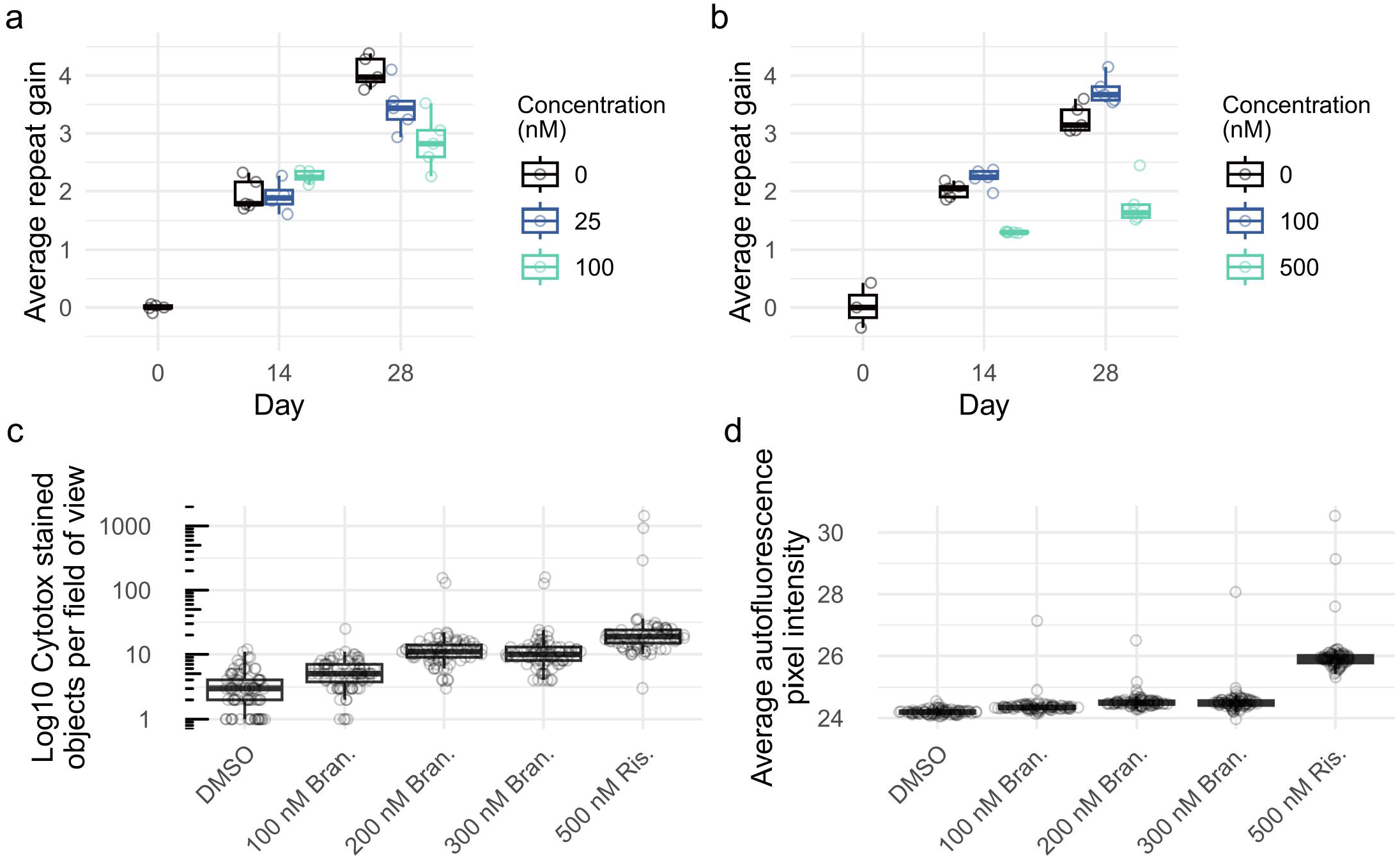
Branaplam and risdiplam treatments reduced repeat expansion in RPE1-AAVS1- CAG115 cells. Average repeat gain of non-induced RPE1-AAVS1-CAG115 cells with treatment of either branaplam (a) or risdiplam (b), with the color indicating the drug concentration. Each treatment group and timepoint had five cultures analyzed, except risdiplam day 0 which had three. (c) Drug cytotoxicity quantified by high-throughput image analysis of cells treated with DNA labeling of dead cells. (d) Average background autofluorescence pixel intensity. For c and d, 81 images were analyzed per treatment.

Increasing drug concentrations are expected to be associated with progressively more potent effects at both target and off-target sites, so we assessed whether the reduction in repeat expansion might also be associated with increasing drug cytotoxicity. From high-throughput image analysis assays, cell proliferation was reduced in a dose-dependent manner beginning at 250 nM branaplam and 500 nM risdiplam (Supplementary figure 5a). In the same experiment, acute cytotoxicity assessed by DNA labeling of dead cells showed a dose-dependent increase starting at 500 nM branaplam, but no increase for risdiplam up to 2000 nM (Supplementary figure 5b). To investigate if the drugs caused cell death longer-term, we maintained the cells at confluency for two weeks. Compared to DMSO treatment, we observed a 3-fold increase in DNA labeling of dead cells for 200 nM branaplam (p <0.001) and a 23-fold increase for 500 nM risdiplam (p <0.001) (Figure 5c). We also observed a rise in the background fluorescence in the 500 nM risdiplam group (Figure 5d, Supplementary figure 6), suggesting drug-induced cellular stress, which has previously been correlated with an increase in autofluorescence ^20^. Thus, the effects on CAG instability at the highest drug doses are accompanied by coincident cytotoxicity, potentially due to increasing off-target effects on splicing at loci across the genome.

### HD genetic modifier PMS1 contains a drug-inducible pseudoexon

We postulated that even for low dose branaplam, the suppression of *HTT* CAG repeat instability was likely an indirect consequence of its splice modulation, either at *HTT* or at another locus. Therefore, we analyzed the list of genes with branaplam- and risdiplam-induced pseudoexons described in the RNAseq results of previously published datasets (Supplementary data 1). Two, *PMS1* and *DHFR*, are within haplotypes associated with genetic modification of HD age-at- onset ^5^. The haplotype at *DHFR* also contains the adjacent *MSH3*, a known modifier of repeat instability, but RNAseq data from Bhattacharyya et al. (Supplementary Data 1) showed that branaplam treatment significantly reduced *DHFR* mRNA but not *MSH3* mRNA ^14^. Consequently, we focused on the huntingtin and PMS1 as potential mediators of the splice modulators’ effects on repeat expansion.

*PMS1* contains a pseudoexon centrally located within the 26 kb or 34 kb intron 5 (Figure 6a), depending on the isoform (Supplementary figure 7). In LCLs and RPE1 cells, the predominant isoform a includes exon 6 and, with drug treatment, the pseudoexon is spliced into mRNA for both isoforms (Supplementary figure 8a). The 91 bp pseudoexon contains a stop codon (Supplementary figure 8b), predicted result in a truncated PMS1 lacking the crucial C-terminal MLH1 dimerization domain and potentially to trigger nonsense-mediated decay ^21^. No surrounding polymorphic variants are predicted to affect splicing, with only very rare variants within 50 bp either side of the pseudoexon (Figure 6b). The drug-binding motif differs in the exon upstream of the 5’ss from the *HTT* pseudoexon, with AAUGA at *PMS1* compared to GCAGA at *HTT*, but both have the same downstream intronic guaag motif. Branaplam was more effective for causing *PMS1* pseudoexon inclusion in LCLs with an IC50 of 100 nM compared to 205 nM for risdiplam (Figure 6c). Consequently, the drugs differ in their relative effects on *HTT* and *PMS1* pseudoexon inclusion: branaplam can preferentially target *HTT* (∼4- fold higher IC50 for *HTT* over *PMS1*), while risdiplam preferentially targets *PMS1* (∼3-fold higher IC50 for *PMS1* over *HTT*).

**Figure 6.**
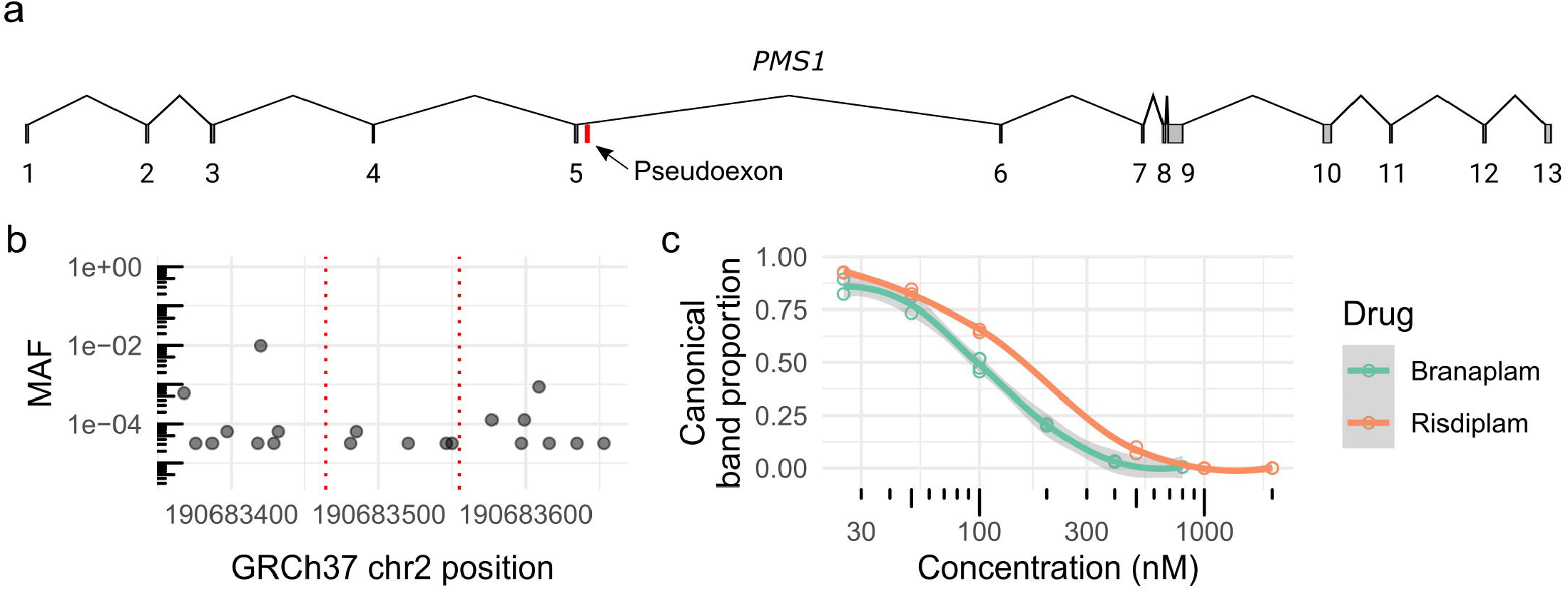
HD modifier *PMS1* contains a drug-inducible pseudoexon. (a) Schematic diagram of the *PMS1* transcript (NM_000534) highlighting the pseudoexon location in red. (b) Minor allele frequency (MAF) of variants 50 bp surrounding the pseudoexon location (red dotted lines). (c) Dose response of *PMS1* exon 5-6 after branaplam (teal) or risdiplam (red) treatment with each empty dot representing a biological replicate and the line showing the local polynomial regression.

### Branaplam suppresses CAG expansion by downregulating PMS1

To determine whether pseudoexon inclusion at *HTT* or *PMS1* was responsible for reducing *HTT* CAG repeat expansion, we edited the pseudoexon locations in these two genes. Using gRNAs directly targeting the GA 3’-exonic motif (Figure 7a, left) at the *HTT* pseudoexon 5’ss, we efficiently generated indels (Supplemental Figure 9). Edited clones had an A insertion between the GA 3’-exonic motif and the GT 5’-intronic motif (Supplemental Figure 10a). In a comparable strategy, attempts with two different gRNAs for the *PMS1* pseudoexon yielded very inefficient editing directly at the site (Supplementary Figure 9). Therefore, we modified *PMS1* with an alternative strategy to delete a 137 bp region from the pseudoexon into the adjacent intron using dual gRNAs (Figure 7a, right). Of the 33 clones isolated, 12 had a heterozygous deletion, but none was biallelically edited (Supplemental Figure 10b).

**Figure 7.**
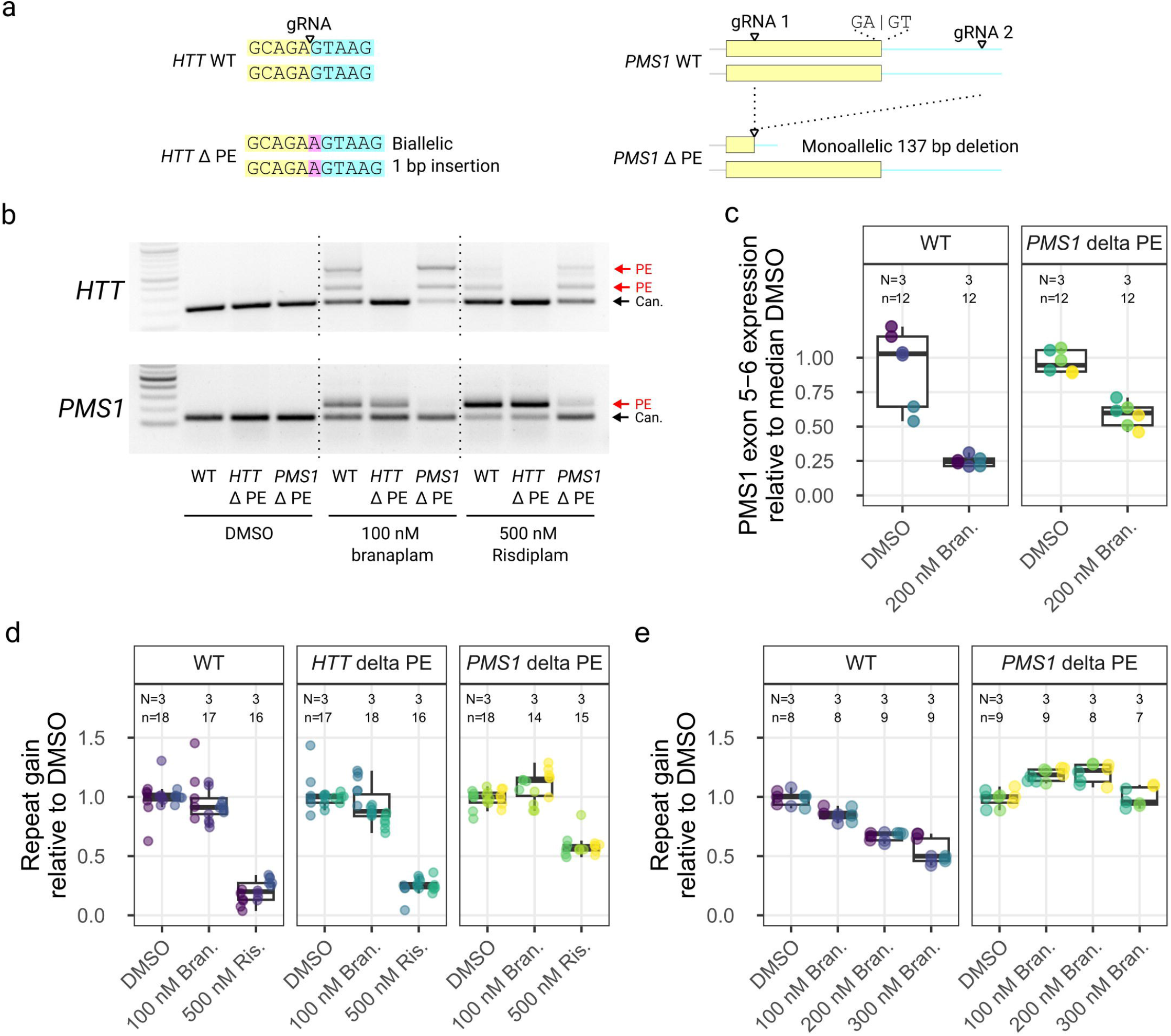
*PMS1* pseudoexon inclusion explained the effect on repeat expansion with branaplam, but only partially with risdiplam. (a) Schematic diagrams showing the CRISPR- Cas9 targeting approach for the disruption of pseudoexon (PE) sequences in *HTT* (left) and *PMS1* (right). Yellow indicates pseudoexon sequence upstream of the 5’ splice site targeted by the drug, blue representing the downstream intronic sequence, with the inserted sequence highlighted in purple. (b) PCR analysis over the *HTT* (top) and *PMS1* (bottom) pseudoexon splice junctions with branaplam or risdiplam treatment for the control and pseudoexon edited cell lines. (c) Accurate quantification of *PMS1* canonical isoform by ddPCR for the control and *PMS1* pseudoexon edited cell lines. The dot color represents a unique cell line. (d, e) The average repeat gain per week after branaplam or risdiplam treatment for the different edited cell lines (dot color), normalized on the average repeat gain in the DMSO for each genotype.

We treated representative *HTT-* and *PMS1-*edited lines with the splice modulators to determine the effect of the genome editing on both canonical and drug-induced splicing. In the former, the A insertion disrupted the drug-induced pseudoexon inclusion, resulting in only canonical splicing from *HTT* exon 49-50 (Figure 7b) despite treatment with 100 nM branaplam or 500 nM risdiplam. In the latter, these treatments markedly increased the proportion of canonical *PMS1* splice product (Figure 7b). Accurate quantification of the *PMS1* canonical isoform by ddPCR showed that the *PMS1* monoallelic editing did not change the level of splicing across the *PMS1* exon 5-6 junction in the absence of drug (p = 0.6 relative to wild-type) (Figure 7c). However, 200 nM branaplam treatment elicited a 3.8-fold (95% CI: 2.8 to 5.7, p < 0.0001) reduction in wild-type cells but only a 1.6 fold (95% CI: 1.4 to 1.9, p < 0.0001) reduction in the *PMS1*-edited cells (p = 0.0003) (Figure 7c). Overall, disrupting the sequences required for *PMS1* pseudoexon inclusion reduced the effectiveness of the splice modulators but did not affect canonical splicing.

We next quantified the repeat instability in these cell lines in 4-5 week experiments with various drug treatments. There were systematic clonal differences in the rate of repeat expansion (Supplementary Figure 11bc), so we normalized the data to the repeat expansion in the DMSO group for each clone. We treated the cell lines with either 100 nM branaplam for relatively stronger *HTT* splice modulation or 500 nM risdiplam for relatively stronger *PMS1* splice modulation. The removal of the *HTT* pseudoexon had no effect on repeat expansion for either 100 nM branaplam or 500 nM risdiplam (Figure 7d), ruling out the drugs’ effects on *HTT* pseudoexon inclusion as the cause of reduced CAG repeat expansion. By contrast, the heterozygous removal of the *PMS1* pseudoexon resulted in weak evidence of a 1.1-fold (95% CI: 0.99 to 1.2, p = 0.019) increase in repeat gain due to 100 nM branaplam treatment (compared to DMSO). With 500 nM risdiplam, the *PMS1* pseudoexon edited cells showed 1.7- fold reduced (95 CI: 1.4 to 2.0, p < 0.0001) repeat gain compared to DMSO, far less than the 4.6-fold reduction (95% CI: 3.5 to 6.7, p < 0.0001) elicited in wild-type cells (Figure 7d), suggesting that pseudoexon inclusion at *PMS1* makes a substantial contribution to risdiplam’s inhibition of CAG expansion at high dosage.

We repeated the experiment with increasing doses of branaplam to confirm the decrease in repeat expansion in wild-type cells and the enhanced repeat expansion in the *PMS1* pseudoexon-edited cells. In wild-type cells, we again observed a dose-dependent effect of branaplam on preventing CAG repeat expansion, which decreased 1.2-fold at 100 nM branaplam (95% CI: 1.1 to 1.2, p < 0.0001), 1.5-fold at 200 nM (95% CI: 1.4 to 1.6, p < 0.0001) and 1.9-fold at 300 nM (95% CI: 1.7 to 2.1, p < 0.0001) relative to DMSO (Figure 7e). By contrast, the *PMS1* pseudoexon-edited cells displayed repeat expansion increased by 1.2-fold at 100 nM (95% CI: 1.1 to 1.3, p < 0.0001) and 200 nM (95%CI: 1.1 to 1.3, p < 0.0001), with 300 nM appearing similar to DMSO (p = 0.62) (Figure 7e). Overall, the results of targeting *PMS1* via the drug inducible pseudoexon explained the reduction in rate of CAG repeat expansion caused by branaplam but only partially explained the observed effect with risdiplam. The partial effect with the latter along with the increases in expansion with the lower branaplam doses, suggest that the drugs may also have effects on splicing in other genes that influence CAG repeat instability.

## Discussion

Orally-available small molecule splice modulators provide an attractive option for therapeutic development, especially for genetic diseases of the nervous system. Their potential has been demonstrated by the Federal Drug Administration (FDA) approval of risdiplam for treatment of spinal muscular atrophy (SMA), where it promotes inclusion of exon 7 in *SMN2,* whose product then compensates for *SMN1*-inactivating mutations. Branaplam was also tested in SMA patients ^22^. Exploration of the genome-wide effects of these and related splice modulators led to the recognition of the pseudoexon in *HTT* and the potential for this mechanism to yield a treatment for HD. Our data indicate that for HD, another therapeutically-relevant splice modulator target is *PMS1*, which has already been validated by human genetics as a modifier of disease onset, providing a fundamentally different alternative to strategies based on reducing mutant huntingtin.

GWAS for modifiers of HD age-at-onset and other clinical landmarks identified *PMS1* among several DNA repair genes also implicated as modifiers of CAG repeat instability, including *FAN1*, a suppressor of repeat expansion ^23,24^, and other mismatch repair genes encoding members of the MutSβ (*MSH3*), MutLα (*MLH1*, *PMS2*), and MutLβ (*MLH1*, *PMS1*) complexes ^5^. Given that *PMS2* and *MLH1* are key genes whose inactivation is a cause of Lynch syndrome ^25^, *MSH3* and *PMS1* appear to be preferable targets for potential therapeutic downregulation in HD. Additionally, in contrast to *Mlh1* and *Pms2*, the loss *Pms1* does not cause tumors in mice. ^26^. While the role of *MSH3* as an enhancer of CAG repeat expansion has been well established ^27–29^, there are few studies on the role of *PMS1* in somatic repeat expansion, perhaps due to its unclear function in canonical human mismatch repair ^30^. The GWAS determined that *PMS1* harbors both clinical landmark-hastening and -delaying variants that are common in the human population, but their mechanism has not been established. However, damaging *PMS1* variants in exome sequencing of HD individuals associated with extremely delayed HD onset suggest that reduced PMS1 function suppresses somatic CAG expansion ^31^. Our demonstration that *PMS1,* like *MSH3,* is required for CAG repeat expansion in a human cell line model strongly supports this conclusion. Loss of PMS1 has also been shown to largely prevent expansion of the CGG repeat in a mouse embryonic stem cell model of the fragile X-related disorders ^32^. Interestingly, in our study and the mouse CGG repeat model, there was a small degree of repeat expansion remaining after knocking out *PMS1*. This is complicated by targeting exon 6 in both cases, which can be spliced out to form a minor *PMS1* isoform whose function remains unclear. The reduction of expansion from inactivating PMS1 in CGG and CAG repeats indicates its broader relevance as a potential target for therapeutic downregulation across repeat disorders. The presence of a modulable pseudoexon in *PMS1* provides a new strategy to achieve its downregulation via small molecules.

*PMS1* knockout heterozygotes had the same repeat expansion characteristics as non-edited cells, indicating that one active PMS1 allele is sufficient to support CAG repeat instability in these cells. However, while branaplam and risdiplam both promote *PMS1* pseudoexon inclusion, albeit with different potency, when one allele was made refractory to splice modulation, the drugs had distinct outcomes with respect to CAG repeat instability. Risdiplam continued to reduce CAG repeat expansion, albeit less robustly, while branaplam increased CAG expansion slightly. Thus, the drugs might also impact on one or more other genes involved with repeat instability. As an example, high risdiplam dosage results in downregulation of another HD genetic modifier, *LIG1* ^33^, suggesting that a deeper exploration of the differential drug effects on CAG repeat expansion in this model might yield additional modifiers and greater mechanistic understanding.

The differential potency and effects highlight the complex nature of these splice modulating drugs with many targets and changes in gene expression. As we have demonstrated, an added layer of complexity is the impact of genetic variation in influencing effects of the drugs at both target and off-target loci. For *HTT*, we identified rare variants that affected pseudoexon inclusion whose impact would depend on the chromosome carrying them. On the non-expanded *HTT* chromosome, the outcome might be positive, allowing continued expression of wild-type huntingtin, whereas on the expanded CAG chromosome, continued expression of mutant but lower expression of wild-type would be more likely to have a deleterious outcome. Another concern with genetic variation is the potential for unexpected off-target effects. We identified many such potential variants, most of which were very rare, but across many individuals, the likelihood of a patient with such a variant receiving drug is non-trivial. Our approach was biased, relying on known branaplam-responsive exons. However, identifying novel pseudoexons activated by genetic variation would be an important next step. Clearly, human genetic variation should be taken into consideration with therapeutics that target specific genetic sequences, whether it involves CRISPR-Cas modification ^34^ or small molecules as described here. Encouragingly, we show that AI tools can be used to identify the genetic variants and therefore potential off targets, which allows an approach of screening patients before they receive such interventions.

While all of the above factors must be considered carefully in developing a potential therapeutic, these small molecule splice modulators have huge delivery advantages with their oral availability and broad distribution, including into the cortex and striatum ^15^. Indeed, inherent in their differential potency and off-target effects is the promise that chemical modifications and a better understanding of the mechanism of splice modulation can identify compounds that more specifically target *PMS1* and reduce potential side-effects. The drugs are proposed to drive alternative splicing by stabilizing non-canonical nGA 3’-exonic motifs at the 5’ss ^14,15^. Our results with the *in vivo* editing of the *HTT* pseudoexon 5’ss support that mechanism, with a single A insertion between the exonic and intronic splice motifs preventing pseudoexon splicing. However, this editing prevented both pseudoexon inclusion (exon 50a) and the generation of the alternative product (exon 50b) that does not use this pseudoexon 5’ss. The exon 50b product was detectable in the RNAseq results of previous publications ^14,35^, but was not focused upon since it results in the same frame-shifting outcome. We speculate that this product can fit within the nGA 3’-exonic motif stabilization model through the order of intron splicing and intron retention, which can be driven by the relative strength of the splice sites ^36^. When we weakened the intron 49 upstream splice site in a minigene, we observed a decreased ratio of exon 50b product relative to the exon 50a product. Additionally, the strong effect of genetic variants near the *HTT* pseudoexon 3’ splice site suggests an important role for this 3’ss region in the drugs’ efficacy. There may also be alternative explanations, with the drugs having an unexplained component to their mechanism. Indeed, a recent publication challenges how branaplam interacts with the U1 / 5’ss, proposing that there are two interaction modes, one for the nGA 3’- exonic motif stabilization and a second interaction with the surrounding sequence ^37^. It also suggests that cocktails of the splice modulators show synergy and can influence the target specificity ^37^. Together with further chemical modification, this synergy increases the options for identifying splice modulating therapeutics that specifically target *PMS1* for repeat expansion disorders and, ultimately, that target other genes in diseases where modulating alternative splicing could prove beneficial. For HD and other CAG repeat disorders, the cell line system that we have developed, which shows significant CAG expansion in confluent cultures, will facilitate the discovery, testing and development of such therapeutic approaches.

## Methods

### LCLs and drug treatment

This work was approved by the Mass General Brigham Institutional Review Board. Lymphoblastoid cell lines (LCLs) were generated from HD patients as previously described ^38^. LCLs were grown in suspension in RPMI 1640 medium (MilliporeSigma, 51536C), with 15% fetal bovine serum (MilliporeSigma, F0926). For branaplam (Synonyms: LMI070, NVS-SM1) (MedChemExpress, HY-19620,) or risdiplam (Synonyms: RG7916; RO7034067) (MedChemExpress, HY-109101) treatments, a 1 mM stock solution prepared in DMSO was diluted in media to the concentrations indicated for 24 hours. Each experiment had the same cell line treated as a control, which was used to correct for run-run variation for the gel-based PCR quantification. LCLs were genotyped by microarray and imputed as previously described ^5^.

### RNA isolation, cDNA synthesis, PCR, and densitometry

RNA was isolated using TRIZOL reagent (Invitrogen, 15596026) following the manufacturers protocol. Any contaminating genomic DNA was removed using ezDNase (Invitrogen, 11766051) following the manufacturers protocol. The cDNA was synthesized using the Superscript IV kit (Invitrogen, 18091050) with poly(A) oligo(dT) with an incubation at 50°C and 80°C for 10 min each, followed by an incubation with RNase H at 37°C for 20 min.

The relative pseudoexon inclusion was quantified by PCR from exons flanking the pseudoexon (Supplementary table 1). We used GoTaq G2 Hot Start PCR kit (Promega, M7423) with the following conditions: initial denaturation 94 °C (2 min), 40 cycles of 94 °C (30 s), 60 °C (30 s), 72 °C (45 s), final extension 72 °C (5 min). Amplicons were loaded onto a 2% agarose gel with EZvision (VWR, 97064-190) and the band intensity was quantified by densitometry using ImageJ ^39^.

### Minigene cloning, mutagenesis, and transfection

A minigene construct was prepared by isolating the entire *HTT* exon 49-50 region of interest (Supplementary table 1) from HEK293T genomic DNA. We used the Q5® High-Fidelity PCR Kit (New England Biolabs, E0555S) with the following conditions: initial denaturation 98 °C (3 min), 35 cycles of 98 °C (10 s), 64 °C (30 s), 72 °C (60 s), final extension 72 °C (2 min). This PCR fragment was TOPO cloned into pcDNA™3.1/V5-His backbone (Invitrogen, V81020). We used *in vivo* assembly cloning ^40,41^ for site directed mutagenesis to modify the nucleotide 1 bp upstream of the exon 49 splice junction to each of the alternative nucleotides (Supplementary table 1). The PCR for cloning was with UltraRun® LongRange PCR Kit (QIAGEN, 206442) with the following conditions: initial denaturation 93 °C (3 min), 18 cycles of 93 °C (30 s), 60 °C (15 s), 68 °C (3 min 35 s), final extension 72 °C (10 min). The amplicons were treated with *Dpn*I restriction enzyme to remove the plasmid template and transformed into XL10 gold competent cells prepared by ‘Mix and Go!’ transformation kit (Zymo Research, T3001). The sequence of the isolated plasmids was confirmed using nanopore sequencing (Plasmidsaurus, SNPsaurus LLC). Confirmed plasmids were transfected into HEK293T cells with lipofectamine 3000 (Invitrogen, L3000001) following the manufacturer’s protocol.

### ddPCR gene expression quantification

Absolute expression quantification was carried out with the QX200 Droplet Digital PCR (ddPCR, Bio-Rad). We used the primer mix for probes (no dUTPs) (Bio-Rad, 1863023) and AutoDG Instrument (Bio-Rad, 1864101) for automated droplet generation following the manufacturer’s instructions. All primers and probes are listed in Supplementary table 1.

### Predicting the effect of genetic variation on pseudoexon splicing

To predict the effect of genetic variation on all known genes with pseudoexons, we used pseudoexons identified from RNAseq in three publications ^14,15,17^ and used a previously described approach ^42^. Briefly, sequences were taken 50 bp either side of each of the pseudoexon splice sites, with *in silico* saturation mutagenesis to modify each position to the other three alternative nucleotides, followed by using spliceAI ^16^ to predict effect of each variant on pseudoexon splicing based on the flanking exons of the gene.

### RPE1-AAVS1-CAG115 model generation

The RPE1-AAVS1-CAG115 model was generated by targeted knock-in of a *HTT* exon1 fragment into the AAVS1 safe harbor locus. We isolated the entire exon 1 of *HTT* with 115 CAG repeats from an HD patient with UltraRun® LongRange PCR Kit (QIAGEN, 206442) with supplementation of 10% DMSO under the following conditions: initial denaturation 93 °C (3 min), 35 cycles of 93 °C (30 s), 61 °C (15 s), 68 °C (60 s), with a final extension of 72 °C (10 min). The primers (Supplementary table 1) had flanking *Sal*I sites which were used to insert the HTT fragment as a GFP fusion-protein (Supplementary Figure 12) in an all-in-one tetracycline-inducible expression cassette with AAVS1 homology arms (AAVS1-TRE3G-EGFP was a gift from Su-Chun Zhang (Addgene plasmid # 52343; http://n2t.net/addgene:52343; RRID:Addgene_52343). This plasmid contains promotor-less puromycin resistance gene with a 3’ splice site that generates puromycin resistance when correctly inserted into intron 1 of *PPP1R12C* (also known as AAVS1) ^43^. hTERT RPE-1 (CRL-4000 - ATCC) were transfected with lipofectamine 3000 (Invitrogen, L3000001) following the manufacturer’s protocol with AAVS1 targeting vector and predesigned transcription activator-like effector nucleases (hAAVS1 TALEN Left and Right were gifts from Su-Chun Zhang, Addgene plasmid # 52341 & 52342; http://n2t.net/addgene:52341; http://n2t.net/addgene:52342; RRID:Addgene_52341; RRID:Addgene_52342). Since hTERT RPE-1 already has expression of puromycin resistance gene, we selected with a high 20 µg/mL dosage of puromycin for 1 week. Clones were isolated by limited dilution and were screened for presence of transgene insertion by PCR of the 5’ homology arm over the puromycin resistance gene (Supplementary table 1). We used GoTaq G2 Hot Start PCR kit (Promega, M7423) with the following conditions: initial denaturation 94 °C (2 min), 35 cycles of 94 °C (30 s), 60 °C (30 s), 72 °C (60 s), final extension 72 °C (5 min).

### Cytotoxicity analysis

Acute cytotoxicity was quantified in RPE1-AAVS1-CAG115 cells with Incucyte SX5 (Sartorius) high throughput image analysis. Cells were seeded at 5000 cells per well and imaged every 2 hours for three days. We also treated with Incucyte® Cytotox Red Dye (Sartorius, 4632) following manufacturer’s instructions. The cell confluency and count of cytotox stained nuclei was quantified using the Incucyte software.

For cytotoxicity in long-term culture, we grew the cells to confluency and treated with selected drug concentrations for two weeks. We treated with Incucyte Cytotox Red Dye and analyzed the cells after 20 hours. We used a custom pipeline to count the number of dead cells as well as quantify the background autofluorescence. For counting dead cells, we set a threshold and segmented stained nuclei using the python scikit-image package ^44^, with a minimum object size of 5 pixels to exclude artifacts. For the autofluorescence analysis, calculated the mean pixel intensity above the background but below the threshold used to identify the stained nuclei.

### Repeat instability analysis

We carried out CAG repeat instability experiments with a high-throughput plate-based pipeline from growing the cells all the way through to capillary electrophoresis. The RPE1-AAVS1- CAG115 were seeded into 96-well plates and grown to confluency to trigger contact inhibition, which enables analysis of repeat expansion in the absence of cell division. The cells were fed every 2-3 days for a total of 4-6 weeks, with genomic DNA isolated using the Quick-DNA 96 Kit (Zymo Research, D3011).

Repeat tracts were quantified by PCR amplification followed by capillary electrophoresis. We used the Taq PCR Core Kit with Q solution (Qiagen, 201225) with 5 µL of the isolated genomic DNA following PCR conditions: initial denaturation 95 °C (5 min), 30 cycles of 95 °C (30 s), 65 °C (30 s), 72 °C (1 min 30 s), final extension 72 °C (10 min). We optimized the PCR with the nested design to only amplify the transgenic exon 1 fragment, which we used for the instability experiments following pseudoexon editing. This PCR had an outer amplicon (Supplementary table 1) for 12 cycles under the same conditions above, followed by the standard fragment analysis assay for the inner amplicon with an additional 22 cycles. Amplicons were analyzed using a 3730XL DNA Analyzer (36 cm array, POP-7 Polymer, standard fragment analysis conditions) with 0.8 ul PCR product is loaded in 9.4 ul Hi-Di Formamide (Applied Biosystems), with 0.1 ul GeneScan 500 LIZ (Applied Biosystems). The fragments were identified and converted to bp sizes using GeneMapper 5.0 (Applied Biosystems). Repeat lengths for each fragment within a sample was calculated from linear models fit using samples with known repeat lengths for each run.

We calculated a repeat instability metric ‘average repeat gain’, describing the average number of repeat units a population of repeat fragments changes from a defined starting point, similar to what was described previously ^45^. We first defined a window of 40 repeat units either side of the identified modal repeat for each sample, with a fragment height threshold of 5% of the modal repeat height. The weighted repeat length was then calculated for each sample by finding the weighted arithmetic mean of the CAG repeat length using the peak height as the weighting. The average repeat gain was the difference between the weighted repeat length for a timepoint and the starting timepoint. When there were multiple timepoints, average repeat gain per week was calculated by fitting a linear modal with a fixed intercept through the average repeat gain at time 0, then finding the slope. With just one timepoint, the average repeat gain was divided by the number of weeks.

### Genome editing

Various CRISPR-Cas9 approaches were used for genome editing experiments in RPE1- AAVS1-CAG115 cells. We used CRISPick ^46,47^ to select gRNAs (Supplementary table 2).

For the HD modifiers we cloned oligos encoding the spacers of the gRNAs into pSpCas9(BB)- 2A-Blast, which was a gift from Ken-Ichi Takemaru (Addgene plasmid # 118055; http://n2t.net/addgene:118055; RRID:Addgene_118055). The plasmids were transfected into RPE1-AAVS1-CAG115 using the 4D-Nucleofector X Unit (Lonza) and the P3 4D-Nucleofector™ X Solution (V4XP-3024) following the manufacturer’s protocol and the EA-104 Nucleofector program. The cells were treated with 25 µg/mL Blasticidin for 4 days, followed by an additional 10 µg/mL for 7 days selection. To amplify *FAN1*, *MSH3*, and *PMS1* (Supplementary table 2) target sites, we used the Q5® High-Fidelity PCR Kit (New England Biolabs, E0555S) with the following conditions: initial denaturation 98 °C (3 min), 35 cycles of 98 °C (10 s), 60 °C (30 s), 72 °C (60 s), final extension 72 °C (2 min). We pooled amplicons from the four different genes together and sequenced with llumina MiSeq via the MGH Center for Computational and Integrative Biology DNA core. CRISPResso pooled ^48^ was used to demultiplex the reads and quantify editing outcomes.

The polyclonal cell populations were found to be edited with 83%, 33%, and 57% indels for *FAN1, MSH3*, and *PMS1*, respectively (Supplementary figure 4a). The most common edits in each population were single bp insertions for *MSH3* (25% of reads) and *PMS1* (43% reads), but for *FAN1*, the most common edit was a 99 bp deletion (16% of reads). These edits resulted in frameshift in 38% *FAN1*, 32% *MSH3*, and 56% *PMS1* of reads. The *FAN1* population had a large number of deletions, with 46% of reads having a >20 bp deletion, compared to an average 0.7% for the other targets. We analyzed the effect of these perturbations in a 6-week repeat instability experiment. The modal repeat lengths for the initial populations were very similar, with 127 repeats for non-targeting control and *FAN1*, 126 for *MLH3* and 125 for *PMS1*.

To analyze the *MSH3* and *PMS1* clonal strains from these edited pooled populations, we genotyped the clones with a barcode multiplexing strategy. Up to eight samples were uniquely barcoded with a unique identifier sequence on the forward primer, with the amplicons pooled, sequenced as described above, demultiplexed *in silico*, and each clone’s read analyzed with CRISPResso. Clones were called homozygous when the top editing outcome accounted for more than 85% of the two most frequent aggregated editing outcomes, otherwise they were called heterozygous.

For precisely targeting the pseudoexon location, we manually selected gRNA sequences with predicted cut sites within 3 bp of the splice site. We cloned oligonucleotides encoding the gRNA spacers into BPK1520 (Addgene plasmid # 65777) to generate gRNA expression plasmids. These plasmids were co-transfected with wild-type SpCas9 (RTW3027, Addgene plasmid # 139987) or the SpG variant capable of targeting sites encoding NGA PAMs (RTW4177, Addgene plasmid # 139988) (Supplementary table 2). The plasmids were transfected with nucleofection as described above and GFP positive cells were FACS sorted with FACSAria™ III Cell Sorter (BD Life Sciences) 48 hours after transfection. The editing was quantified by Sanger sequencing trace decomposition ^49^ and confirmed by sanger sequencing ion the isolated clonal strains by Sanger sequencing. For *PMS1* deletion of pseudoexon, two gRNAs flanking the 5’ pseudoexon splice site were transfected as described above with the pSpCas9(BB)-2A-Blast vector. Clonal cell strains were screened for deletion by PCR with primers flanking the *PMS1* pseudoexon location (Supplementary table 1).

### Statistics

The data were analyzed with R ^50^ and the tidyverse suite of packages ^51^, and marginaleffects ^52^. P-values are the result of two-tailed t-tests. All data, graphs and statistics are available with executable R code (https://github.com/zachariahmclean/2023_splice_modulators).

## Author contributions

ZLM: Conceptualization, Methodology, Software, Formal analysis, Investigation, Data Curation, Writing - Original Draft, Visualization, Project administration. DG: Software and Formal analysis prediction of variants on splicing. KC: Software for repeat instability, cytotoxicity image analysis, genotyping, and phasing. JCLR: Methodology modifier CRISPR-Cas9 development. SS: Methodology and Software for modifier CRISPR-Cas9 development sample genotyping. INF: Investigation Figure 3. ZENVM: Investigation Figure 2. MR: Software cytotoxicity analysis. EM: Resources minigene cloning vector, Critical Reading. JR: Experimentation cell culture, Resources LCLs. TG: Experimentation CAG sizing, sequencing. DL: Resources human subjects. BPK: Methodology and Resources CRISPR-Cas9 pseudoexon editing Figure 7. JML: Resources identification of LCLs. MEM: Resources identification of LCLs, Supervision, Critical Reading. VCW: Conceptualization, Resources. RMP: Conceptualization, Resources, Methodology modifier CRISPR-Cas9 development and sample genotyping. JFG: Conceptualization, Resources, Writing - Original Draft, Supervision, Project administration, Funding acquisition.

## Acknowledgements

Supported by Hereditary Disease Foundation Fellowships (Z.L.M. and J.C.L.R.), NIH grants NS091161 (J.F.G.), NS126420 (R.M.P.), NS049206 (V.C.W.), NS105709 (J-M.L.), NS119471 (J-M.L.) and DP2-CA281401 (B.P.K.), an MGH ECOR Howard M. Goodman Fellowship (B.P.K.), the CHDI Foundation (J.F.G., M.E.M.), and the Huntington’s Disease Society of America Human Biology Project (R.M.P.). These HD studies would not be possible without the vital contribution of the research participants and their families.

## Competing interests

J.F.G. and V.C.W. were founding scientific advisory board members with a financial interest in Triplet Therapeutics Inc. Their financial interests were reviewed and are managed by Massachusetts General Hospital (MGH) and Mass General Brigham (MGB) in accordance with their conflict of interest policies.

J.F.G. consults for Transine Therapeutics, Inc. and has previously provided paid consulting services to Wave Therapeutics USA Inc., Biogen Inc. and Pfizer Inc.

V.C.W. is a scientific advisory board member of LoQus23 Therapeutics Ltd. and has provided paid consulting services to Acadia Pharmaceuticals Inc., Alnylam Inc., Biogen Inc. and Passage Bio. R.M.P. and V.C.W. have received research support from Pfizer Inc.

B.P.K. is a consultant for EcoR1 capital and Curie.Bio, and is an advisor to Acrigen Biosciences, Life Edit Therapeutics and Prime Medicine. B.P.K. has a financial interest in Prime Medicine, Inc., a company developing therapeutic CRISPR-Cas technologies for gene editing. B.P.K.’s interests were reviewed and are managed by MGH and MGB in accordance with their conflict-of-interest policies.

J-M.L. consults for Life Edit Therapeutics and serves on the scientific advisory board of GenEdit Inc.

E.M. is inventor on an International Patent Application Number PCT/US2021/012103, assigned to Massachusetts General Hospital and PTC Therapeutics entitled “RNA Splicing Modulation” related to use of BPN-15477 in modulating splicing.

## Description of Additional Supplementary Files

File Name: Supplementary Data 1

Description: Branaplam-responsive exons from Monteys et al., 2021 (Extended Data Table 1 & 2); Bhattacharyya et al. (Supplementary Data 2, HTT-C2), 2021; and Keller et al., 2022 (Supplementary Data Table 2). A combined table of each drug responsive exon, the gene, the type (pseudoexon vs existing annotated exon) and the GRCh37/hg19 coordinates.

File Name: Supplementary Data 2

Description: SpliceAI predictions for the effect of variants on the splicing of branaplam responsive exons. The variant coordinates are GRCh37/hg19 position. In the exclusive_events column, TRUE indicates that the branaplam-responsive exon is a pseudoexon while FALSE indicates an existing annotated exon.

## Data availability

All data, graphs and statistics are available with executable R code (https://github.com/zachariahmclean/2023_splice_modulators).

---

Supplementary figure 1. Sanger sequencing for TA-cloned PCR fragments for: (a) *HTT* canonical exon 49-50 splicing, (b) pseudoexon inclusion exon 49-50a-50, (c) alternative splice site exon 49-50b. Yellow is exon 49 sequence, orange is exon 50a, blue is the entire intronic sequence between exon 50a and exon 50, green is exon 50.

Supplementary figure 2. Genetic variants and *HTT* splice modulation. Canonical *HTT* exon 49- 50 for cell lines containing variants rs79689511 (orange) and rs772437678 (teal) for 100 and 200 nM branaplam.

Supplementary figure 3. Branaplam dose response for *TENT2* (left) and *ZFP82* (right) for canonical splice product after branaplam treatment.

Supplementary figure 4. RPE1-AAVS1-CAG115 HD modifier genome editing. (a) Proportion of edited reads in samples transfected with either non targeting or targeting gRNAs in RPE1- AAVS1-CAG115. The numbers above the bars indicate the total reads analyzed. (b) Representative fragment analysis traces of isolated clonal edited strains from either *MSH3* (red) or *PMS1* (orange) targeted populations.

Supplementary figure 5. The effect of branaplam and risdiplam on cell growth and acute cytotoxicity. (a) High-throughput image analysis for quantification of confluency over time with the treatment of branaplam (left) or risdiplam (right). The drug concentration is represented by the color, with key concentrations labelled on the plot. (b) Quantification of DNA labelling of dead cells.

Supplementary figure 6. The effect of selected branaplam and risdiplam concentration on cells treated for two weeks at confluency. Images are shown with no adjustment, highlighting the brightly stained dead nuclei, or with a brightness adjustment to highlight the background autofluorescence.

Supplementary figure 7. *PMS1* alternatively splice variants. NCBI RefSeq curated transcripts for each PMS1 isoform. The pseudoexon location is shown with the dashed red line.

Supplementary figure 8. *PMS1* pseudoexon. (a) PCR from *PMS1* exon 5-7 showing the two variants, isoform a (includes exon 6) or isoform b (skips exon 6) in LCLs or RPE1 in DMSO control cells and the formation of pseudoexon products (red label) for both isoform a and b across increasing branaplam concentrations in LCLs. (b) Sanger sequencing traces from an isolated band of pseudoexon inclusion with a PCR from *PMS1* exon 5-6 PCR, with the termination codon indicated by a star.

Supplementary figure 9. Editing outcomes for *HTT* and *PMS1* direct pseudoexon editing by sanger sequencing and quantified by Sanger sequencing trace decomposition. *PMS1* pseudoexon gRNA 1 also targets an intergenic region on chromosome 13, but editing was not quantified.

Supplementary figure 10. Edited cell clones for *HTT* and *PMS1* pseudoexon disruption. (a) sanger sequencing for *HTT* edited clones. (b) PCR of PMS1 genomic region surrounding pseudoexon location of several isolated cell lines, with the monoallelic deletion highlighted for the three selected cell clones.

Supplementary figure 11. Pseudoexon clone phenotypes. The average repeat gain per week after branaplam or risdiplam treatment for the different edited cell lines (dot color), Figure 7de without DMSO normalization.

Supplementary figure 12. AAVS1-CAG115 plasmid. The AAVS1-TRE3G-EGFP (Addgene plasmid # 52343) plasmid was modified to insert the *HTT* exon 1 coding sequence using *Sal*I restriction sites. The *HTT* exon 1 fragment had an expanded CAG repeat tract with 115 units and was inserted to make a GFP fusion protein using the *Sal*I site as a linker sequence. Annotations of the translation and key motifs are under the appropriate sequences, with the original Addgene plasmid # 52343 sequences indicated in gray.

