## Supplementary information for "*PMS1* as a target for splice modulation to prevent somatic CAG repeat expansion in Huntington’s disease"

Supplementary Figure 1

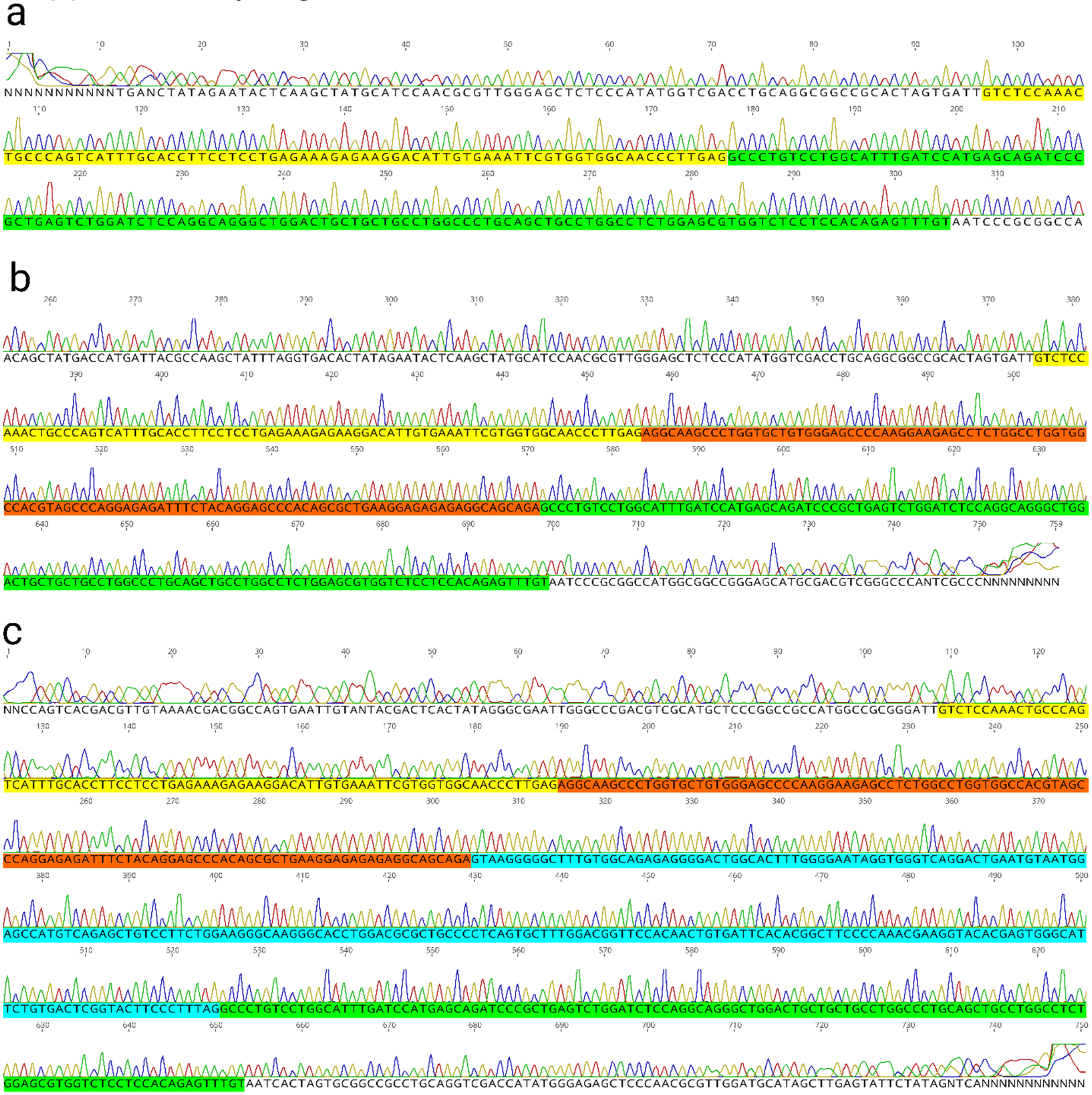

### Supplementary Figure 2

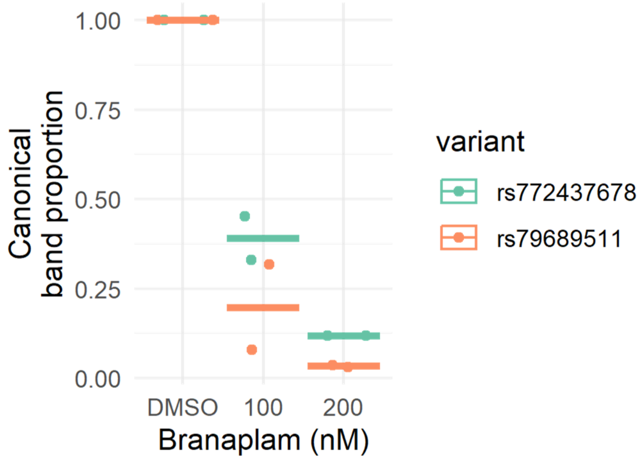

### Supplementary Figure 3

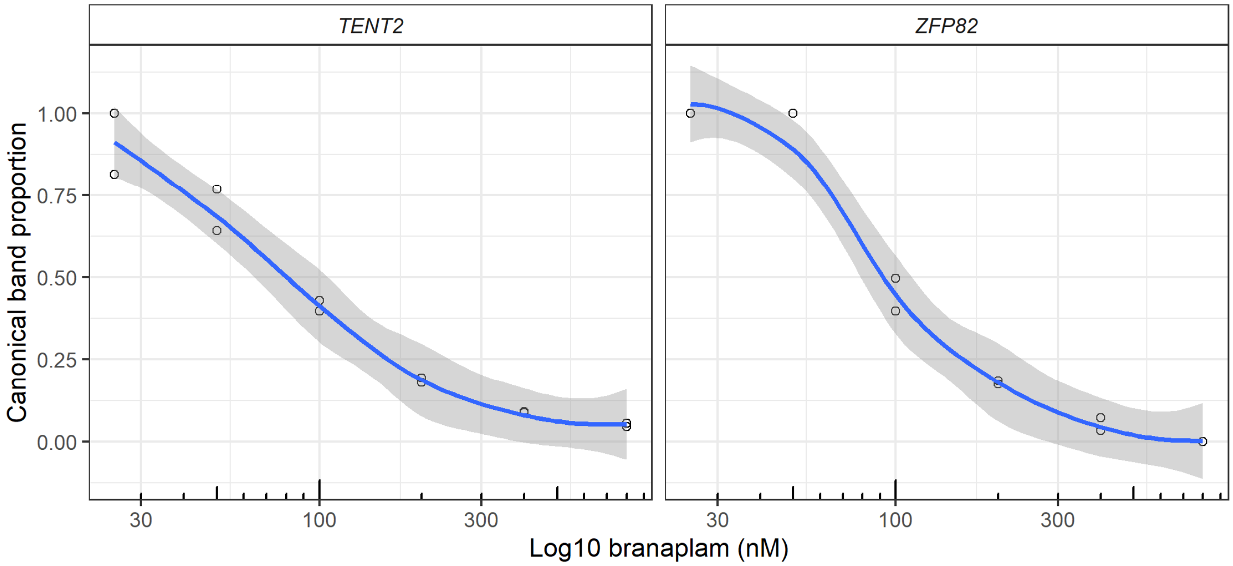

### Supplementary Figure 4

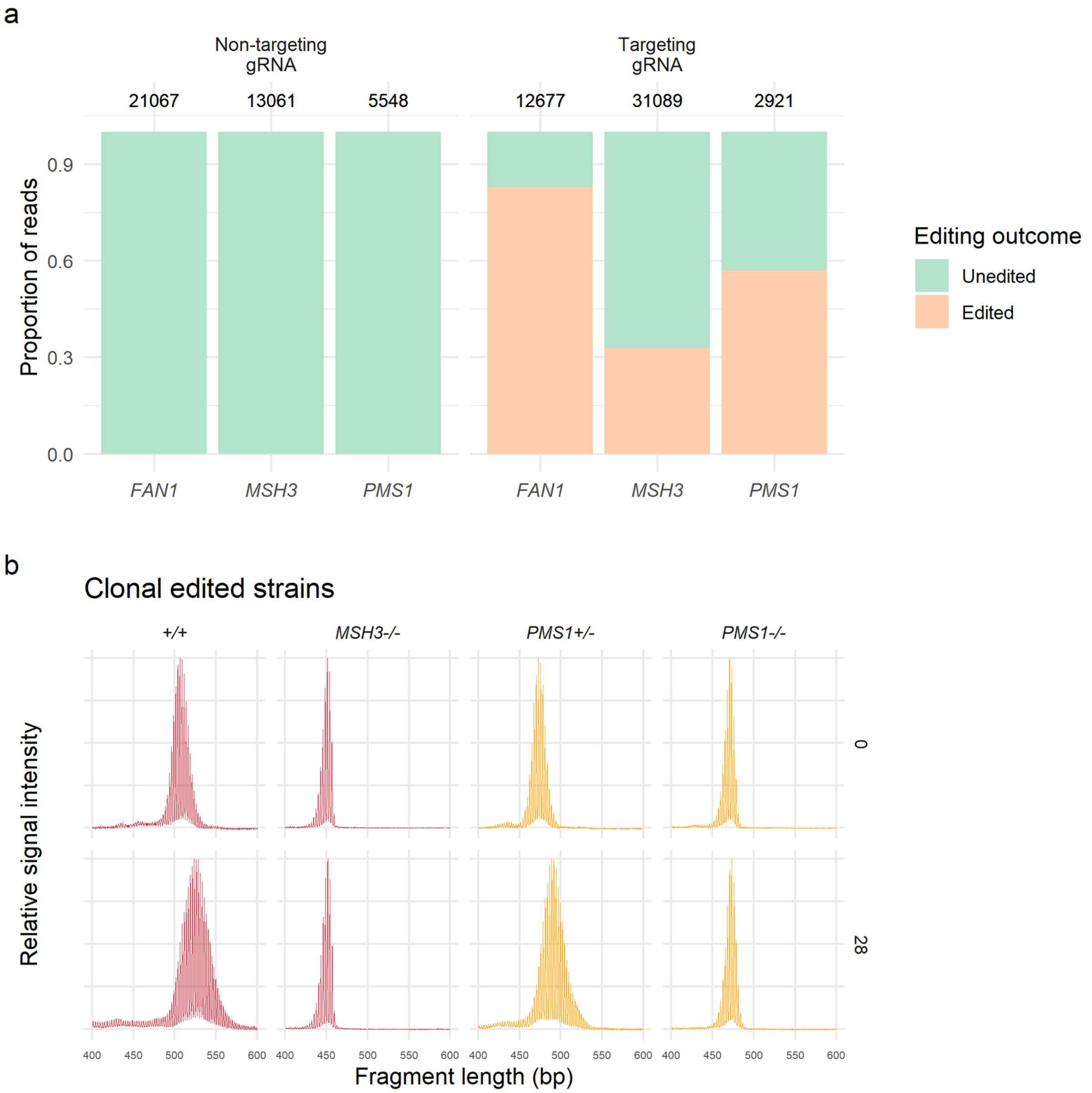

### Supplementary Figure 5

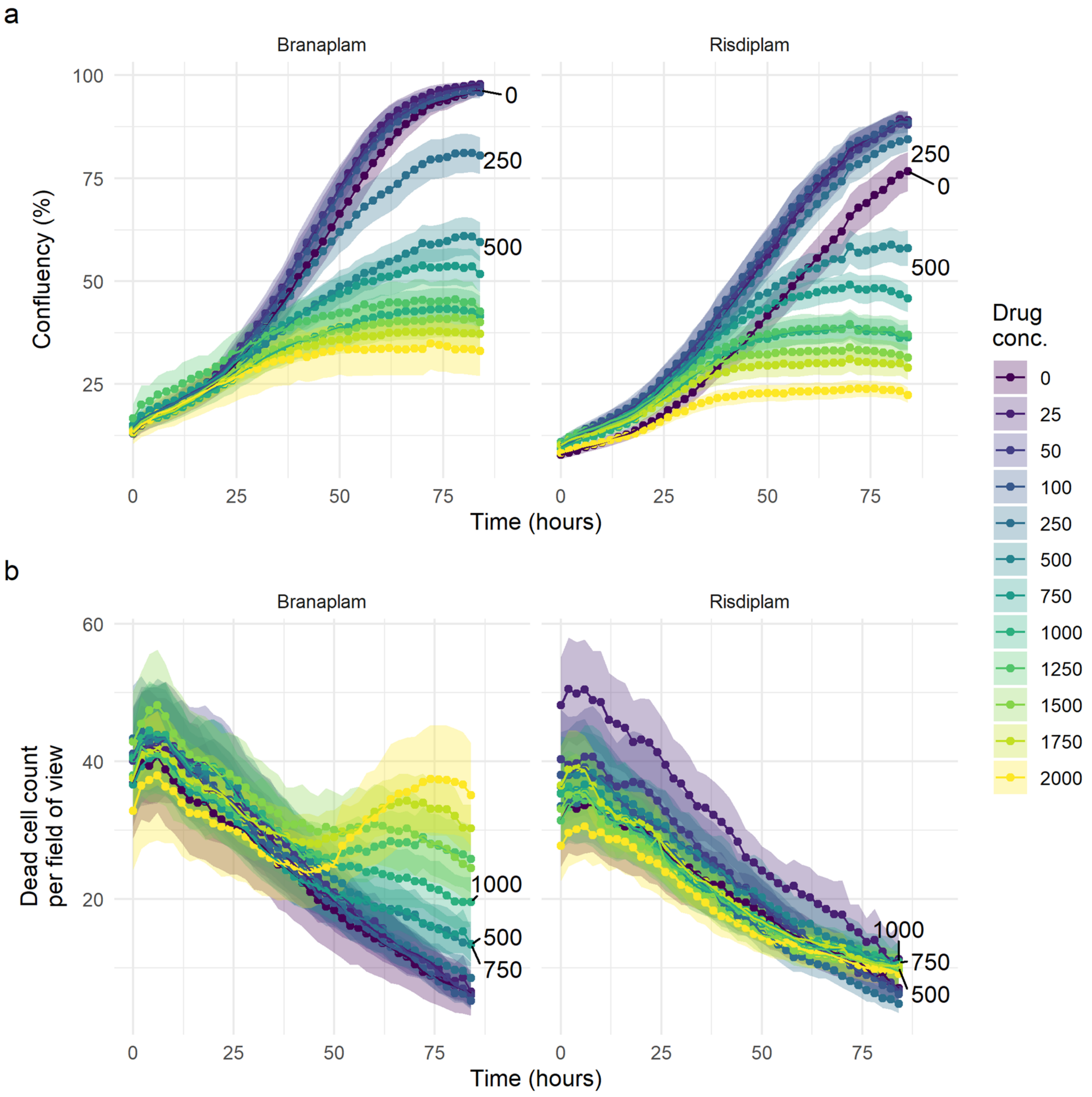

Supplementary Figure 6

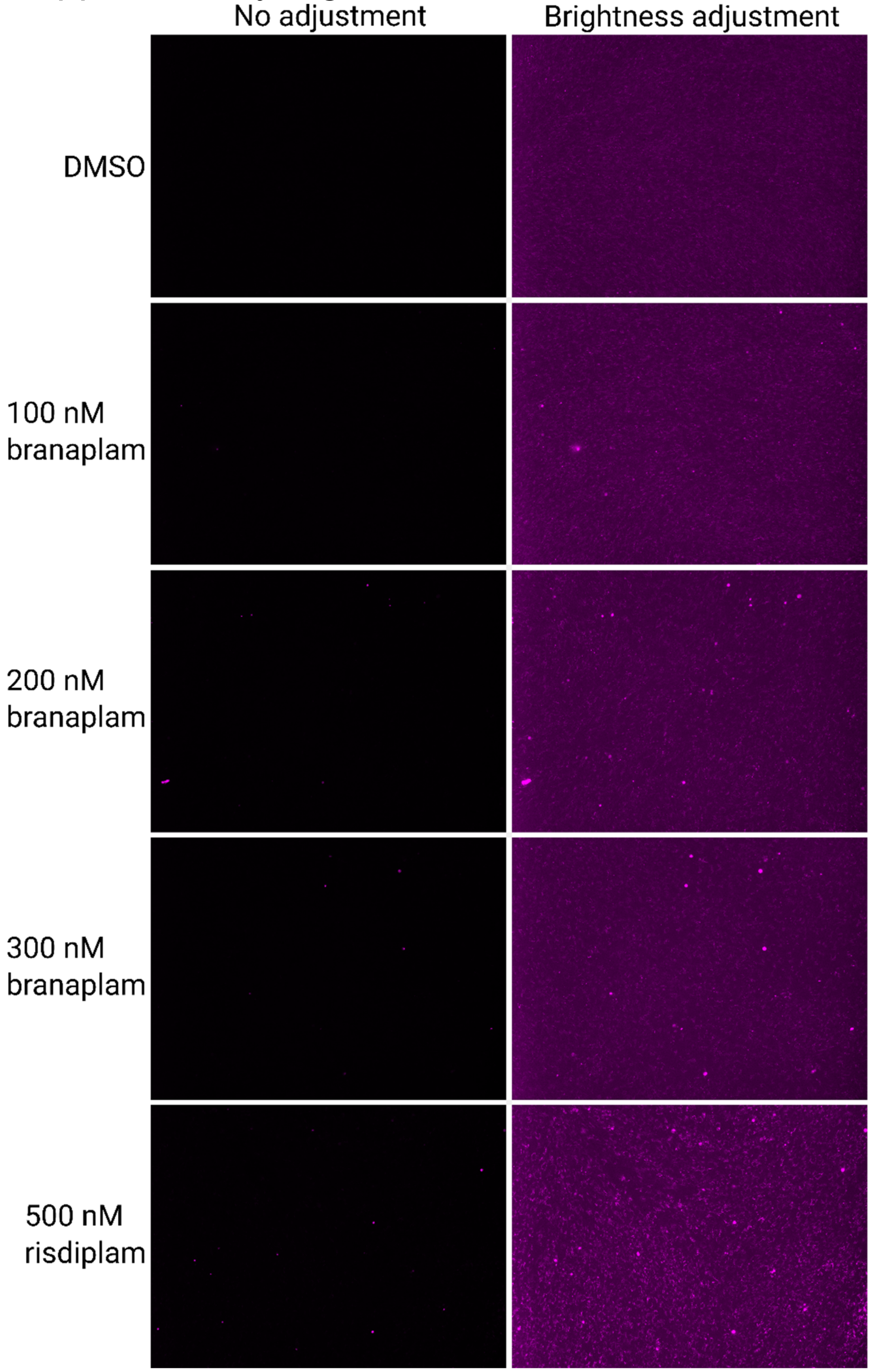

### Supplementary Figure 7

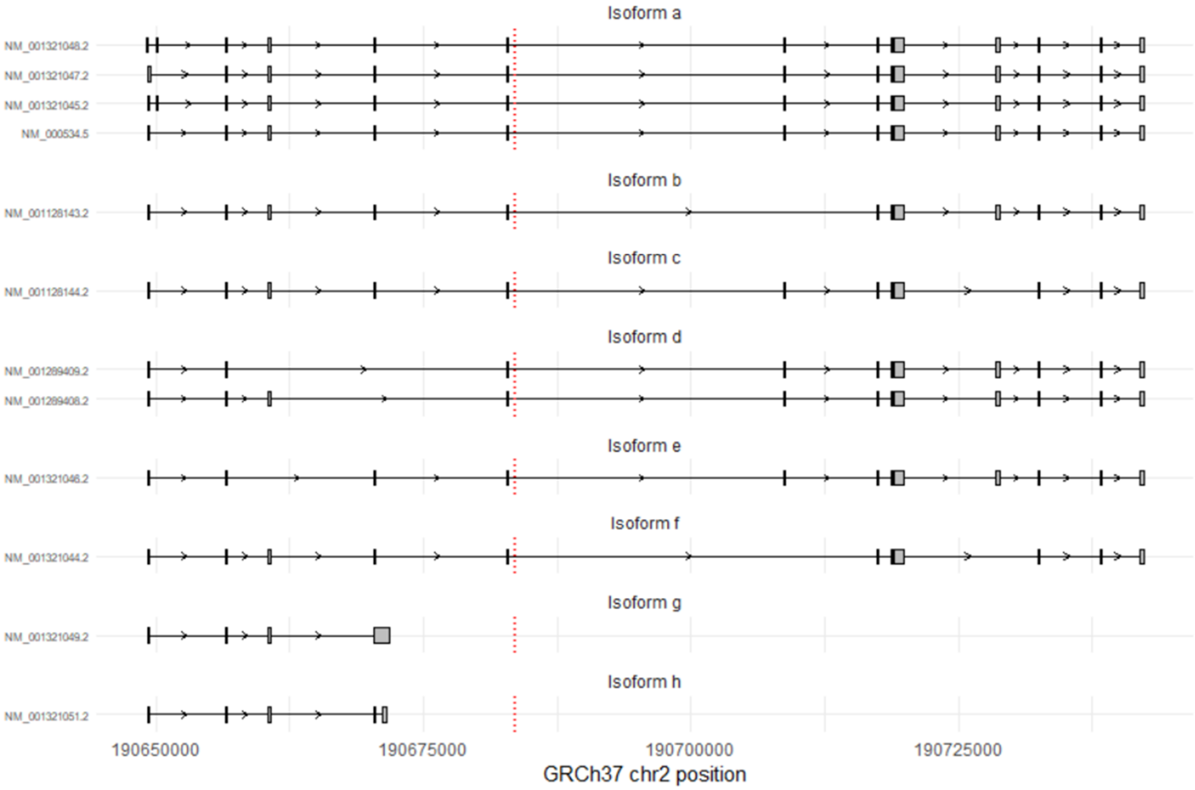

### Supplementary Figure 8

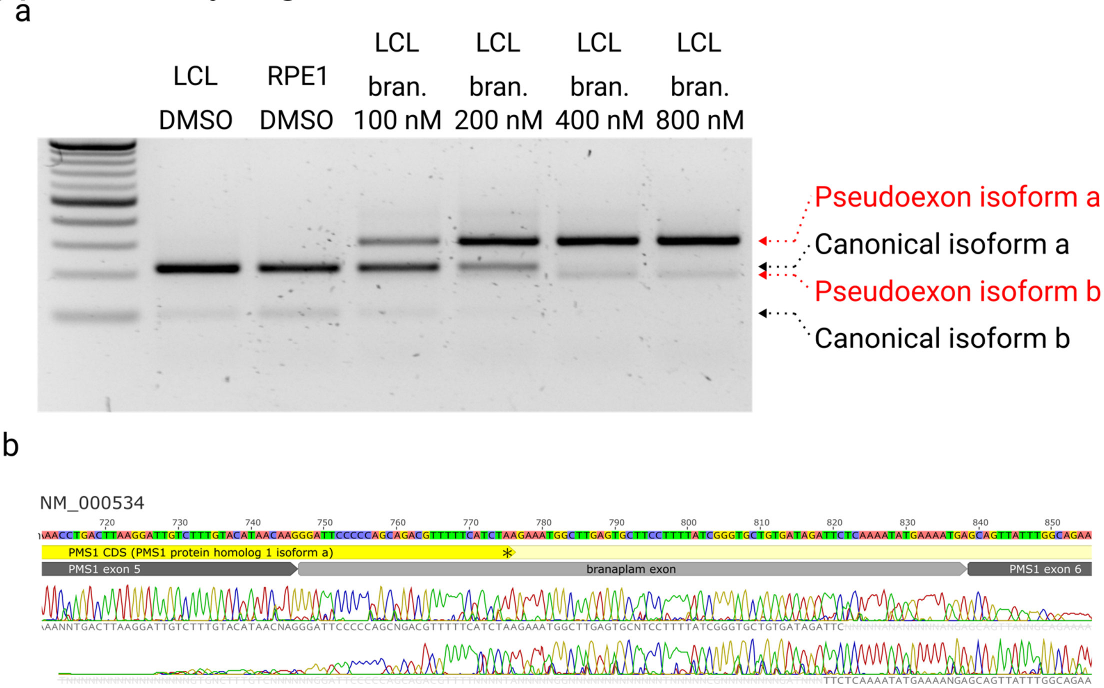

### Supplementary Figure 9

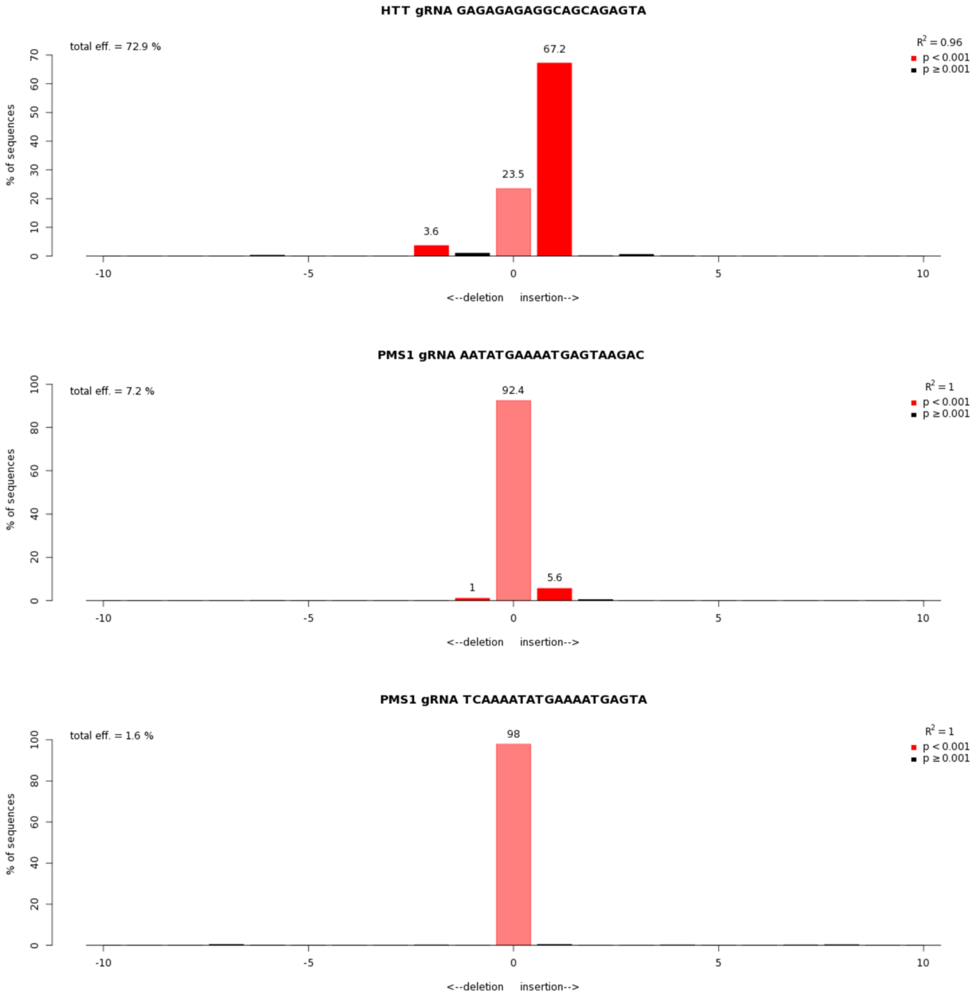

#### Supplementary Figure 10

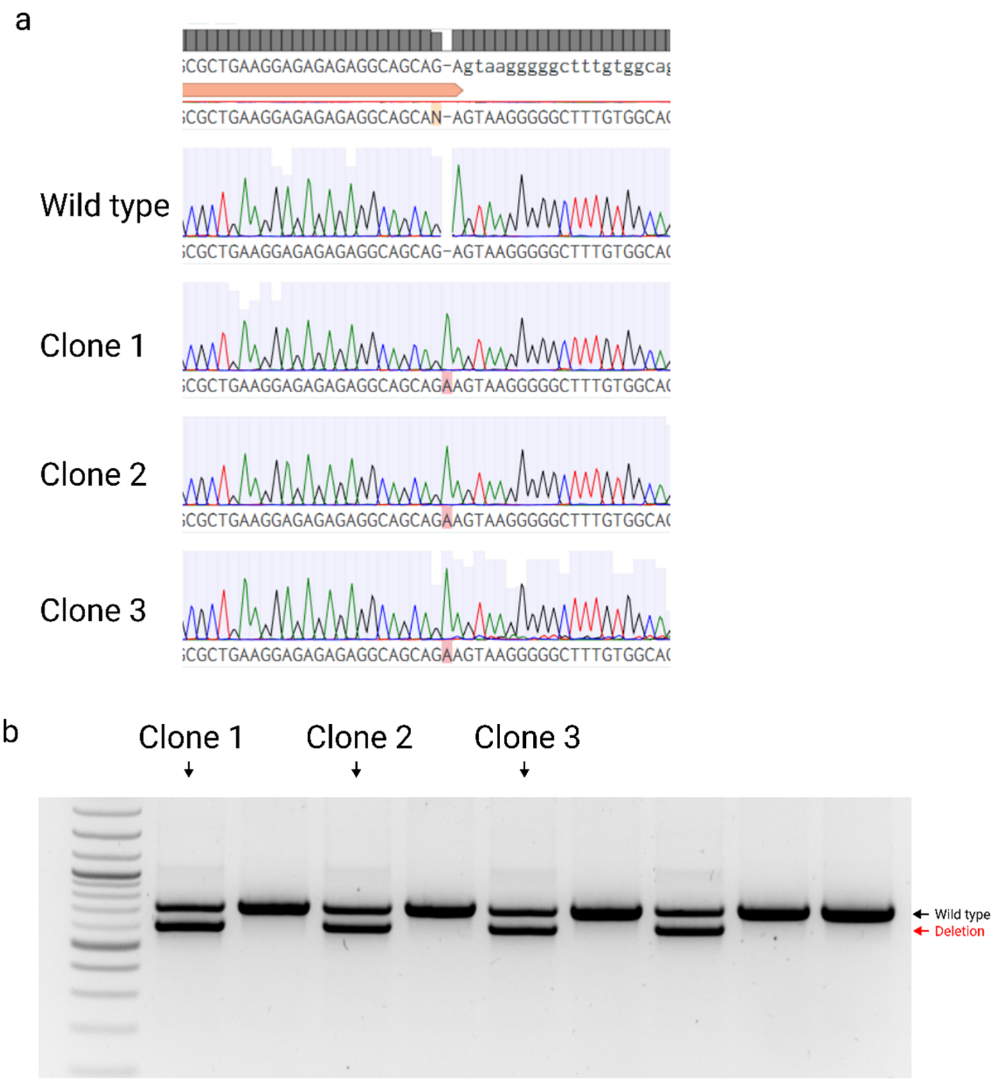

### Supplementary Figure 11

b

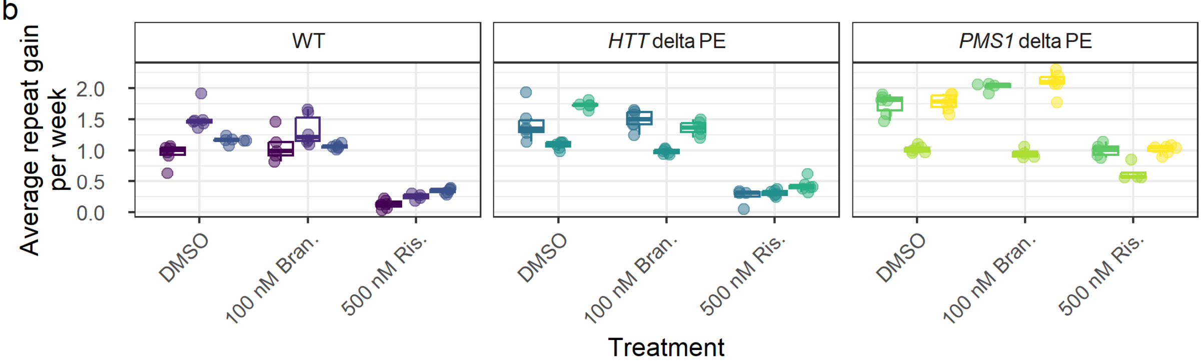

c

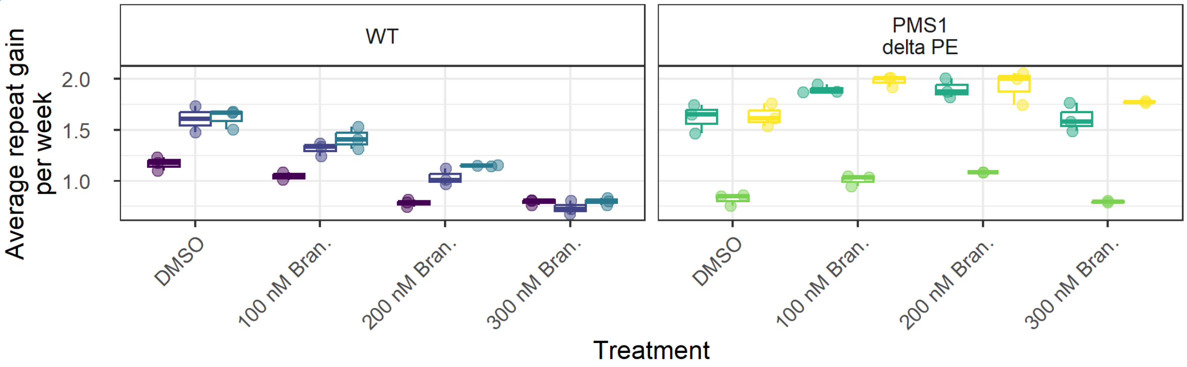

### Supplementary Figure 12

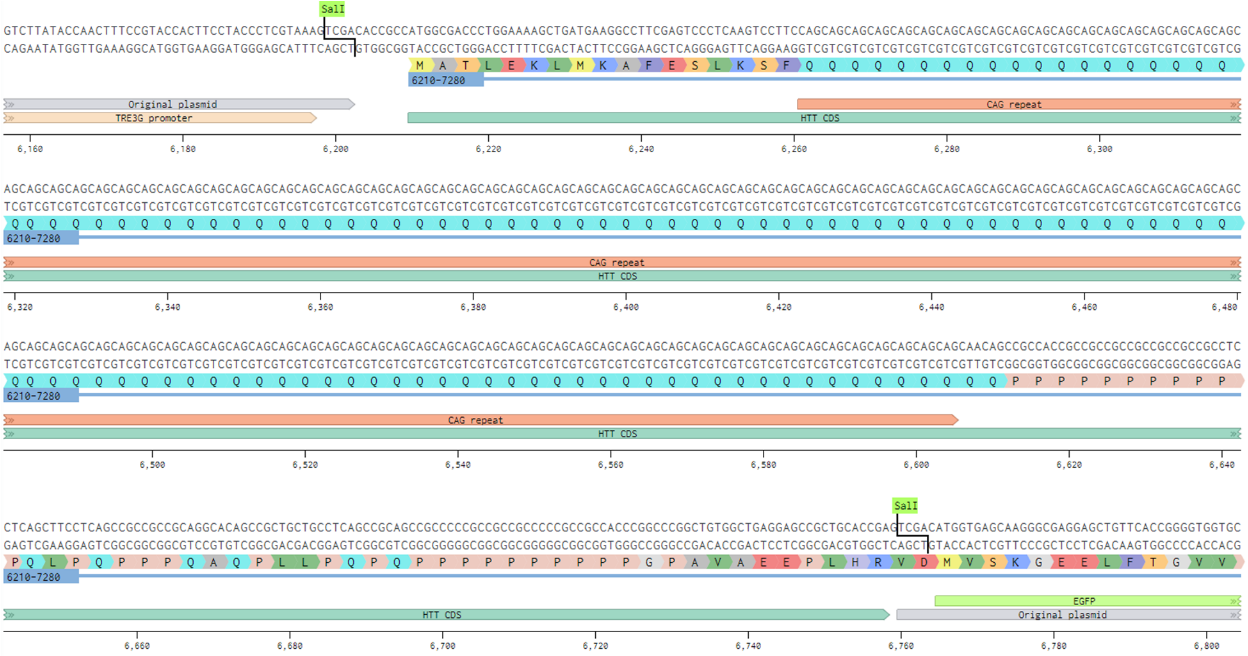

### Supplementary Table 1: primers

| Name | sequence |
|---|---|
| *HTT* exon 49 F | GTCTCCAAACTGCCCAGTCAT |
| *HTT* exon 50 R | ACAAACTCTGTGGAGGAGACC |
| *HTT* exon 49-50 probe | /56-FAM/AACCCTTGAG/ZEN/GCCCTGTCCT/3IABkFQ/ |
| *SDHA* F | TTTGATGCAGTGGTGGTAGG |
| *SDHA* R | CAGAGCAGCATTGATTCCTC |
| *SDHA* probe | /56-FAM/AG CCT AAG A/ZEN/T GAG AGT TCA AGT TGA GTT TGG /3IABkFQ/ |
| *HTT* exon 49 cloning F | ACCATGGGGGATGCTGCACTGTATCAG |
| *HTT* exon 49 cloning R | GGCCTCCAGGATGAAGTG |
| Minigene SDM -1G>C F | GGCAACCCTTGACgtaagaggcagctcgggag |
| Minigene SDM -1G>C R | gctgcctcttacGTCAAGGGTTGCCACCAC |
| Minigene SDM -1G>A F | GGCAACCCTTGAAgtaagaggcagctcgggag |
| Minigene SDM -1G>A R | gctgcctcttacTTCAAGGGTTGCCACCAC |
| Minigene SDM -1G>T F | GGCAACCCTTGATgtaagaggcagctcgggag |
| Minigene SDM -1G>T R | gctgcctcttacATCAAGGGTTGCCACCAC |
| *HTT* exon 1 cloning primers with SalI F | CATGTACGgtcgacaccgccATGGCGACCCTGGAAAAGCTG |
| *HTT* exon 1 cloning primers with SalI R | CATGTACGgtcgacTCGGTGCAGCGGCTCCTC |
| AAVS1 5' homology arm F | CTGCCGTCTCTCTCCTGAGT |
| AAVS1 5' homology arm R | GTGGGCTTGTACTCGGTCAT |
| CAG repeat sizing 1st PCR F (CAG1) | ATGAAGGCCTTCGAGTCCCTCAAGTCCTTC |
| CAG repeat sizing 1st PCR R (EGFP R) | GTCCAGCTCGACCAGGATG |
| CAG repeat sizing 1st PCR F (CAG1) | /56-FAM/ATGAAGGCCTTCGAGTCCCTCAAGTCCTTC |
| CAG repeat sizing 1st PCR R (Hu3) | GGCGGCTGAGGAAGCTGAGGA |
| *FAN1* KO NGS F | AGTGGTTGAAAAACGTGAGGCA |
| *FAN1* KO NGS R | CCATTGCAGCTTGACCCCTGCT |
| *MSH3* KO NGS F | ATGCCCGGCTTGATGCTGTATCG |
| *MSH3* KO NGS R | AGTGTCCTCAAGCTGAAGAACACTGT |
| *PMS1* KO NGS F | GCTGAGGATGAATGCAAAATATAGGA |
| *PMS1* KO NGS R | TCAGAGTGGTACTGAAAGGATTCCA |
| *PMS1* exon5 F | TCTCCTCATGAGCTTTGGTATCC |
| *PMS1* exon6 R | ACAGCAGTCCCCAGAACTGA |
| *PMS1* ex5-6 probe | /56-FAM/TG TAC ATA A/ZEN/C AAG GCA GTT ATT TGG CAG A/3IABkFQ/ |
| *PMS1* exon7 R | TGGTCTGCATCACACTTTGGA |
| *PMS1* int 5 F | AGGCACCACCTATGAACCAG |
| *PMS1* int 5 R | AGCAACAAGGATGGAAATGG |

### Supplementary Table 2: sgRNAs

| Target | sgRNA seq | PAM |
|---|---|---|
| *FAN1* | TGCATGGAGTAACATCCAAG | NGG |
| *MSH3* | AAATTGCCCGACATAGAGAG | NGG |
| *PMS1* | CAGAGTATCAGATCACAAGA | NGG |
| *HTT* pseudoexon 1 | GAGAGAGAGGCAGCAGAGTA | NGG |
| *PMS1* pseudoexon 1 | AATATGAAAATGAGTAAGAC | NGG |
| *PMS1* pseudoexon 2 | TCAAAATATGAAAATGAGTA | NGA |
| *PMS1* pseudoexon flanking LHS | TTAGATGAAAAACGTCTGCT | NGG |
| *PMS1* pseudoexon flanking RHS | CAAAGAATCAACAGTGTAGA | NGG |
